# Cdt1 inhibits CMG helicase in early S phase to separate origin licensing from DNA synthesis

**DOI:** 10.1101/2021.07.06.451311

**Authors:** Nalin Ratnayeke, Mingyu Chung, Tobias Meyer

## Abstract

A fundamental concept in eukaryotic DNA replication is the temporal separation of G1 origin licensing from S phase origin firing. Re-replication and genome instability ensue if licensing occurs after DNA synthesis has started. In humans and other vertebrates, the E3 ubiquitin ligase CRL4^Cdt2^ starts to degrade the licensing factor Cdt1 after origins fire, raising the question of how cells prevent re-replication in early S phase. Here, using quantitative microscopy, we show that Cdt1 inhibits DNA synthesis during an overlap period when cells fire origins while Cdt1 is still present. Cdt1 inhibits DNA synthesis by suppressing CMG helicase progression at replication forks through the MCM-binding domain of Cdt1, and DNA synthesis commences once Cdt1 is degraded. Thus, instead of separating licensing from firing to prevent re-replication in early S phase, cells separate licensing from DNA synthesis through Cdt1-mediated inhibition of CMG helicase after firing.

**Highlights:**

– Cdt1 is present together with fired origins of replication at the start of S phase
– Cdt1 delays DNA synthesis by inhibiting CMG helicase progression after origins fire
– Cdt1 inhibits CMG helicase progression through the MCM-binding domain of Cdt1

## Introduction

In order to duplicate their genome precisely once, eukaryotic cells divide DNA replication into two stages, origin licensing and origin firing. During licensing in G1 phase, cells demarcate future sites of DNA synthesis by loading inactive MCM2-7 helicases onto origins of replication. At the start of S phase, cells begin origin firing, whereby replication factors are recruited to the inactive helicases to form the active CMG (CDC45-MCM2-7-GINS) helicase and replication fork that duplicate DNA (Arias and Walter, 2007; Diffley, 2011; Limas and Cook, 2019). Critically, it is thought that origin licensing must be strictly separated in time from origin firing to avoid re-replication, which occurs when synthesized DNA is re-licensed and replicated again within the same cell cycle (Arias and Walter, 2007; Limas and Cook, 2019; Reusswig and Pfander, 2019). Avoiding re-replication is crucial for maintaining genome stability, and failure to do so results in gene amplification, DNA damage, oncogenesis, and cell death (Arias and Walter, 2007; Pozo and Cook, 2016).

The G1/S transition is a particularly vulnerable period in the cell cycle when cells must simultaneously inactivate licensing activity and initiate origin firing. In humans and other vertebrates, avoidance of re-replication is critically dependent on the repression of the essential licensing factor Cdt1 from the start of S phase through anaphase (Pozo and Cook, 2016). Activation of Cdt1 during this period is sufficient to trigger re-replication (Arias and Walter, 2005a; Dorn et al., 2009; Klotz-Noack et al., 2012; Vaziri et al., 2003), indicating that the presence of active Cdt1 together with synthesized DNA during early S phase could produce re-replication. Cdt1 is repressed by degradation mediated by cullin-RING E3 ubiquitin ligases CRL4^Cdt2^ and SCF^Skp2^ (also known as CRL1^Skp2^), as well as by Geminin binding and hyperphosphorylation by Cyclin A-CDK1, both of which prevent Cdt1 licensing activity (Pozo and Cook, 2016; Zhou et al., 2020). Both Geminin and Cyclin A are degraded during G1 by E3 ubiquitin ligase APC/C^Cdh1^ and only begin to accumulate at the start of S phase (Bastians et al., 1999; Geley et al., 2001; McGarry and Kirschner, 1998), while SCF^Skp2^-mediated Cdt1 degradation does not start until mid-S phase (Grant et al., 2018; Sakaue-Sawano et al., 2017). These findings suggest that CRL4^Cdt2^ alone is responsible for degrading Cdt1 and preventing re-replication in early S phase.

However, the exclusive role of CRL4^Cdt2^ in inactivating Cdt1 at the start of S phase poses a conundrum; for CRL4^Cdt2^ to ubiquitinate and degrade Cdt1 in S phase, Cdt1 must first bind to the replication fork component PCNA (Havens and Walter, 2009, 2011), and therefore Cdt1 degradation can only start after origins have already fired. This regulation would result in a predicted overlap period in early S phase when cells fire origins and could still license DNA before Cdt1 is fully degraded (Arias and Walter, 2007; Havens and Walter, 2011; Reusswig and Pfander, 2019). Since it is expected that fired origins immediately synthesize DNA, this overlap period would be susceptible to re-licensing and re-replication.

Human cells replicate DNA at thousands of sites simultaneously, each of which provides an opportunity for re-replication. Human diploid cells contain approximately 6 gigabases of DNA and typically have an S phase that is 6-10 h long (Cappell et al., 2016; Grant et al., 2018), corresponding to an average rate of DNA synthesis of 10-15 megabases per minute. Thus, even during a short overlap of origin licensing with origin firing, tens to hundreds of megabases of synthesized DNA could be produced. With these considerations in mind, we set out to study the predicted overlap period between firing and licensing in early S phase to understand how cells can protect themselves from re-replication.

Here, using a single-cell microscopy-based analysis of human cells, we show that there is an overlap period in early S phase that lasts approximately 30 min, during which Cdt1 is present together with fired origins in the absence of Geminin and Cyclin A. Strikingly, we show that in addition to licensing origins in G1, Cdt1 has an unexpected second role of inhibiting CMG helicase progression at replication forks during this overlap. This inhibition is dependent on the MCM-binding domain of Cdt1 and is only relieved once Cdt1 is fully degraded or inhibited by Geminin. By delaying DNA synthesis at fired origins during early S phase, cells reduce the amount of synthesized DNA produced in the presence of Cdt1 to deter re-replication. Cdt1-mediated suppression of DNA synthesis fills a critical gap in licensing regulation and allows for uninterrupted protection against re-replication from the first fired origin at the start of S phase to anaphase. Conceptually, our study suggests that instead of temporally separating licensing and firing of origins in early S phase, human cells safeguard genome integrity by using Cdt1-mediated CMG helicase inhibition to separate licensing and DNA synthesis.

## Results

### Cdt1 is present together with fired origins in early S phase

To determine if and for how long Cdt1 is present together with fired origins of replication (Figure 1A), we monitored the degradation of a doxycycline (Dox)-inducible Cdt1-mCherry fusion protein in live MCF-10A cells (a non-transformed human epithelial cell line). To monitor DNA replication, we co-expressed an EYFP-tagged PCNA that forms foci at sites of origin firing and DNA synthesis (Hahn et al., 2009; Leonhardt et al., 2000). In line with previous studies (Grant et al., 2018; Pozo et al., 2018), Cdt1-mCherry degradation at S phase start is coupled to the formation of PCNA foci (Figure 1B, S1A). Time-lapse analysis shows that it takes approximately 30 min between the start and completion of Cdt1-mCherry degradation (Figure S1A), suggesting that there is an extended period in early S phase when Cdt1-mCherry is present together with fired origins.

**Figure 1.**
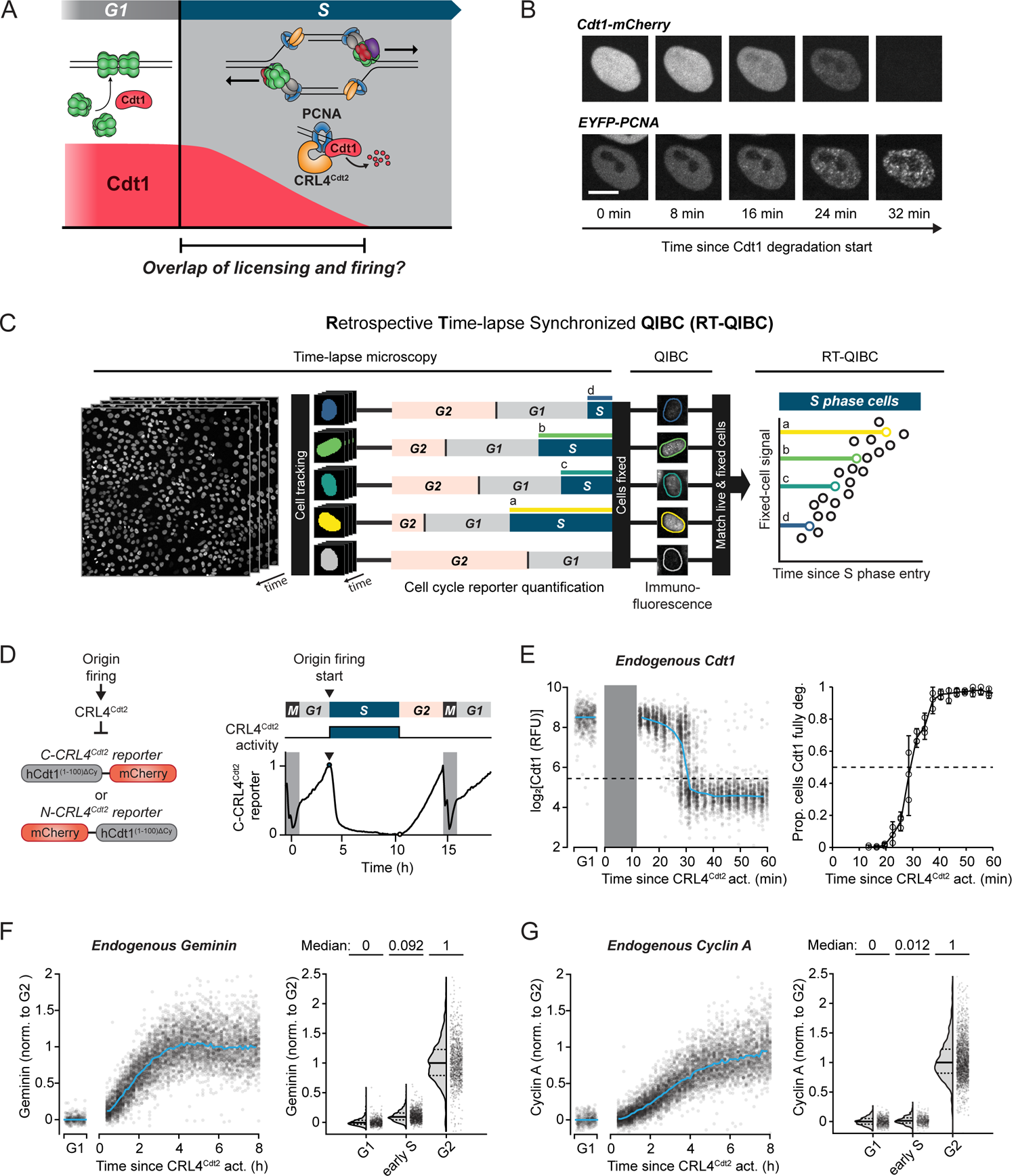
Cdt1 is present together with fired origins in early S phase. **(A)** Predicted overlap between licensing factor Cdt1 and origin firing in early S phase, as Cdt1 is being degraded by CRL4^Cdt2^. **(B)** MCF-10A cells expressing EYFP-PCNA and doxycycline-inducible Cdt1-mCherry (induced 6 h prior to imaging) were imaged using confocal microscopy. Representative of n = 54 cells. Scale bar = 10 µm. Quantification in Figure S1A. **(C)** Diagram of Retrospective Time-lapse Synchronized QIBC (RT-QIBC). **(D)** Left: Live-cell fluorescent reporters of CRL4^Cdt2^ activity. Right: Example single cell trace of C-CRL4^Cdt2^ reporter intensity. Reporter is degraded at S phase start and stabilized at the end of S phase. **(E-G)** RT-QIBC of endogenous protein immunofluorescence (IF) aligned to CRL4^Cdt2^ activation (C-CRL4^Cdt2^ reporter). Cells were live-imaged every 3 min (E) or 8 min (F, G). G1 cells: 1-2 h after anaphase. Dashed and solid lines in violin plots are IQR and median, respectively. **(E)** Left: Cdt1 IF (n = 3,710 S phase cells, 500 G1 cells). Solid curve is median value, dashed line is threshold for fully degraded Cdt1. Grey bar is time period that is not observed due to the requirement of 12 min of reporter degradation to identify S phase start. Representative of 3 independent experiments. Right: Quantification of left. Proportion of cells with Cdt1 levels below detection limit over time within 3 min bins for 3 independent experiments (n = 3,710, 1,873, 1,208 cells, ≥36 cells per bin). Error bars are 95% confidence intervals. **(F)** Left: Geminin IF (n = 13,262 S phase cells, 300 G1 cells). Right: Comparison of Geminin levels in G1 (n = 600 cells), early S (first 30 min, n = 1,051 cells) and G2 (4N DNA and EdU(-), n = 1,063 cells) cells. Representative of 3 independent experiments. **(G)** Left: Cyclin A2 IF (n=13,262 S phase cells, 300 G1 cells). Pool of 10 wells, measured in same cells as Figure 1F (left panel). Right: Comparison of Cyclin A levels from left panel. G1 (n=699 cells), early S (first 30 min, n=503 cells) and G2 (4N DNA and EdU(-), n=2,637 cells) cells. See also Figure S1.

To determine whether endogenous Cdt1 is degraded over a similar time window in early S phase, we utilized a combined quantitative image-based cytometry (QIBC) (Toledo et al., 2013) and live-cell imaging approach (Cappell et al., 2016; Spencer et al., 2013). In this method, we identified the S phase entry time for each cell using automated live-cell imaging of fluorescent cell cycle reporters prior to cell fixation, and identified the same cells in QIBC analysis (Cappell et al., 2016, 2018; Gookin et al., 2017; Stallaert et al., 2021). This method allowed us to retrospectively synchronize fixed-cell measurements of thousands of cells based on the elapsed time from S phase start with high temporal resolution (Figure 1C). For simplicity, we refer to this technique here as Retrospective Time-lapse Synchronized QIBC (RT-QIBC).

To precisely measure S phase entry in live cells, we imaged a reporter of CRL4^Cdt2^ activity based on amino acids 1-100 of human Cdt1, which is rapidly degraded at S phase start by CRL4^Cdt2^ in response to origin firing and reaccumulates at the start of G2 (Sakaue-Sawano et al., 2017) (Figure 1D, S1B). This reporter is not degraded by SCF^Skp2^. We used this reporter in its original N-terminal mCherry-tagged orientation (referred to here as N-CRL4^Cdt2^ reporter), and additionally created and used a C-terminally tagged reporter (C-CRL4^Cdt2^ reporter), which is degraded with slightly faster kinetics at S phase start for precise measurement of the initial moments of S phase (see Methods for discussion of reporters, Figures S1B and S1C). We define the start of S phase to be the start of origin firing and loading of PCNA to replication forks, which triggers CRL4^Cdt2^ activation (referred to in the Figures as CRL4^Cdt2^ act.) and the degradation of the CRL4^Cdt2^ reporters. In line with this, RT-QIBC analysis indicates that the start of degradation of the CRL4^Cdt2^ reporters coincides with chromatin-bound PCNA immunofluorescence staining (Figure S1D).

We performed RT-QIBC of endogenous Cdt1 immunofluorescence staining and aligned asynchronously cycling cells to S phase start. Based on this analysis, endogenous Cdt1 takes approximately 30 min to degrade following the start of origin firing (Figure 1E), similar to the time measured using overexpressed Cdt1-mCherry.

Although Cdt1 is present in early S phase, it could be in an inhibited state either through binding by Geminin or hyperphosphorylation by Cyclin A-CDK1. The levels of Geminin and Cyclin A are expected to be low since they are both degraded by APC/C^Cdh1^ throughout G1 and should only begin to accumulate following APC/C^Cdh1^ inactivation (Cappell et al., 2016; Geley et al., 2001; Limas and Cook, 2019; McGarry and Kirschner, 1998). Using RT-QIBC analysis, we indeed find very low levels of both Geminin (median is 9.2% of G2 levels) and Cyclin A (median is 1.2% of G2 levels) in the first 30 min of S phase, with both gradually increasing following S phase entry (Figures 1F, 1G and S1E). This result is consistent with studies that showed that Geminin and Cyclin A contribute to Cdt1 inhibition later in S and G2 after they accumulate to high enough levels (Klotz-Noack et al., 2012; Zhou et al., 2020).

RT-QIBC analysis was corroborated with live-cell imaging of a reporter of APC/C activity that shows that APC/C^Cdh1^ inactivation (referred to in the Figures as APC/C^Cdh1^ inact.), which is necessary to stabilize Geminin and Cyclin A at the G1/S transition, occurs very near the start of S phase and can occur after the start of S phase (Figure S1B and S1C), in line with previous findings (Grant et al., 2018; Sakaue-Sawano et al., 2017). We conclude that early S phase is characterized by an approximately 30 min-long overlap period, during which replication origins have fired and Cdt1 is still present and active. This presents a problem in the regulation of origin licensing, as synthesized DNA at these fired origins would be susceptible to re-licensing by Cdt1 and re-replication.

### DNA synthesis is inhibited in the presence of Cdt1

We next determined how much DNA is synthesized during the overlap period when origins have fired and Cdt1 is still present. We measured the levels of Cdt1 together with DNA synthesis rates, measured by the incorporation of the thymidine analog 5-Ethynyl-2’-deoxyuridine (EdU) into synthesized DNA in an 8 min period just before cell fixation. Strikingly, as cells transition from G1 to S phase, Cdt1 and EdU staining are mutually exclusive (Figure 2A), arguing that while origins are firing in the presence of Cdt1, there is no detectable DNA synthesis occurring during the overlap period.

**Figure 2.**
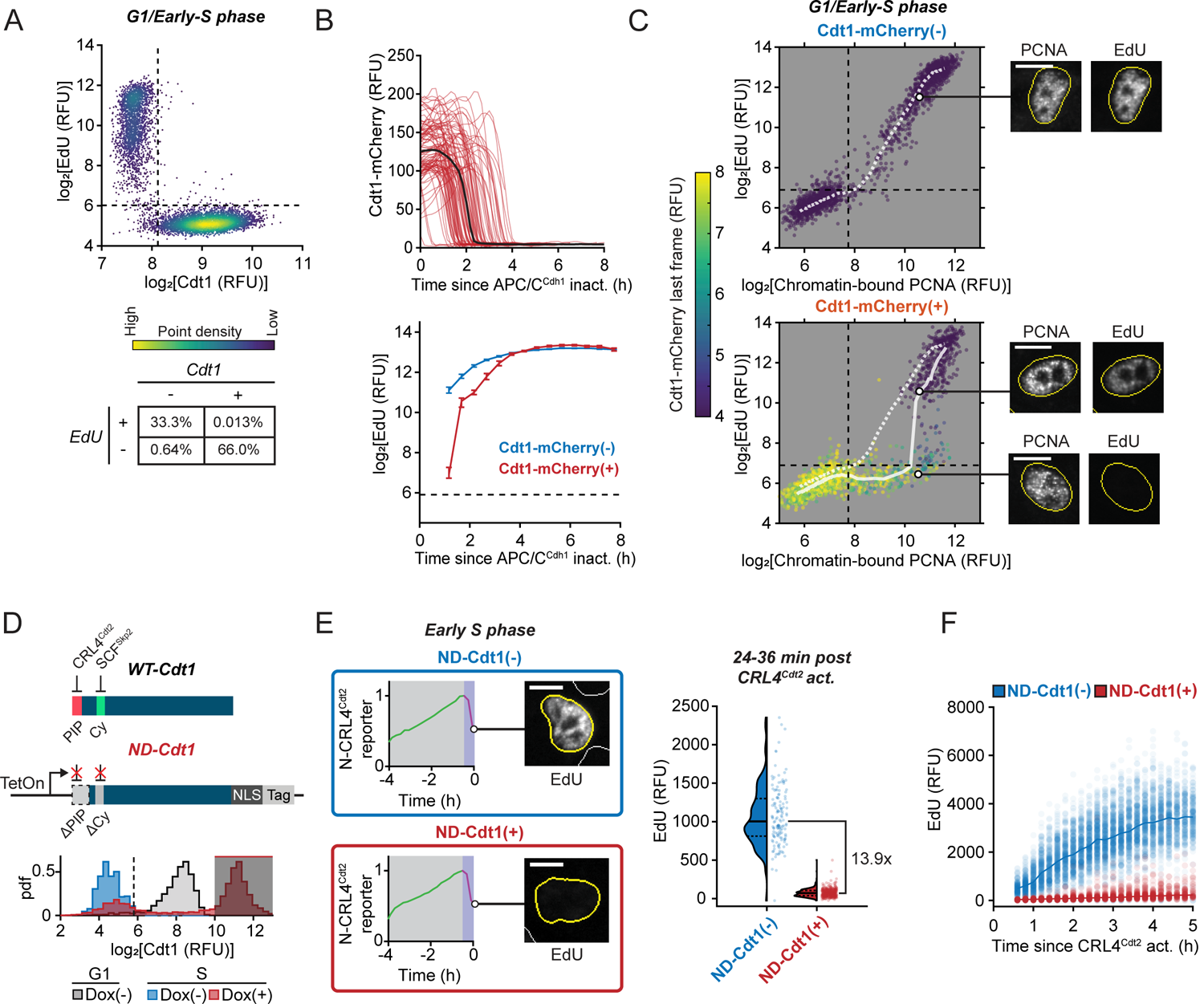
DNA synthesis is inhibited in the presence of Cdt1. **(A)** Quantification of Cdt1 immunofluorescence (IF) and EdU incorporation in cells in late G1 to early S phase (defined by 2N DNA and intermediate Cyclin E/A-CDK activity of 0.7-1.2, see Figure S5A). n = 7,486 cells, representative of 2 independent experiments. Percentage of cells in each quadrant is quantified in bottom table. **(B-C)** RT-QIBC of EdU incorporation and chromatin-bound PCNA in mitogen-released cells with the APC/C reporter and doxycycline (Dox)-inducible Cdt1-mCherry, fixed after 18 h. Cdt1-mCherry(+) cells were identified from Dox-treated cells while Cdt1-mCherry(-) cells were DMSO-treated. Dynamics of mitogen-released cells are in Figures S2A and S2B, and Cdt1-mCherry expression compared to endogenous Cdt1 is in Figure S2C. **(B)** Top: Cdt1-mCherry quantification. Line is median trace (n = 100 traces visualized, median of 7,059 cells). Bottom: EdU incorporation in S phase cells (APC/C^Cdh1^ inactivated with chromatin-bound PCNA). Cdt1-mCherry(-): n=18,367 cells, Cdt1-mCherry(+): 6,970 cells. Line is mean and error bar is bootstrapped 95% confidence interval (n ≥ 93 per bin). **(C)** Chromatin-bound PCNA and EdU incorporation in 2N DNA cells (G1/early S), colored by live-imaged Cdt1-mCherry just prior to pre-extraction and fixation. Cdt1-mCherry(-): n= 3,000 cells, Cdt1-mCherry(+): 2,000 cells. Line is median EdU within bins of chromatin-bound PCNA. Representative cells are shown (scale bar = 10 µm). **(D, E)** RT-QIBC of EdU incorporation in cells overexpressing Dox-inducible non-degradable Cdt1 (ND-Cdt1), induced with Dox and live-imaged N-CRL4^Cdt2^ reporter for 6 h. **(D)** Top: ND-Cdt1 was generated from wild-type Cdt1 (WT-Cdt1) by deleting aa1-19 (removing PIP degron) and mutating the Cy motif (aa68-70), which is necessary for SCF^Skp2^ degradation, to alanine. The mutant was fused to an SV40 NLS to ensure proper localization and either an HA or mCherry tag. Bottom: ND-Cdt1 expression in S phase compared to endogenous Cdt1 by Cdt1 IF. Comparing Cdt1 in G1 cells (1-2 h after mitosis) (grey, n = 2,191 cells), or Cdt1 in S phase cells (0.5-1 h after CRL4^Cdt2^ activation) with ND-Cdt1 induced (red, n = 6,389 cells) or not induced (blue, n = 783 cells). Shaded area represents gate for ND-Cdt1(+) expression selected for EdU quantification in Figure 2E. **(E)** Left: Representative N-CRL4^Cdt2^ reporter traces and EdU stain. 10 µm scale bar. Right: Quantification of cells 24-36 min after CRL4^Cdt2^ activation that divided within 1 h of Dox addition. Dashed and solid lines in violin plots are IQR and median, respectively. ND-Cdt1(-): n = 141 cells, ND-Cdt1(+): n = 400 cells. Representative of 2 independent experiments. **(F)** RT-QIBC of EdU incorporation in mitogen-released cells with the N-CRL4^Cdt2^ reporter, fixed after 18 h. Cells were treated with control siRNA and induced with Dox, performed in same experiment as Figure 3D. ND-Cdt1(+) cells were induced with Dox and selected for expression based on gating in Figure S2F. ND-Cdt1(-): n= 5,500 cells, ND-Cdt1(+): n = 2,000 cells. Line is median value at each time point. Representative of 3 independent experiments. See also Figure S2.

One possible explanation of the lack of EdU incorporation was that Cdt1 itself suppresses DNA synthesis. To explore this possibility, we examined EdU incorporation by RT-QIBC in mitogen-released cells expressing the APC/C reporter together with high levels of Dox-inducible Cdt1-mCherry (Figures S2A-C). Markedly, these cells exhibited a delay in the start of EdU incorporation following APC/C^Cdh1^ inactivation, and this delay closely corresponded to the time during which Cdt1-mCherry was still being degraded (Figure 2B). In line with this interpretation, we identified a prominent population of cells with chromatin-bound PCNA but low EdU incorporation, corresponding to cells that had fired origins but had not yet fully degraded their Cdt1-mCherry (Figure 2C).

Since Cdt1-mCherry was still degraded in S phase, we more directly tested for a suppressive role of Cdt1 by engineering a non-degradable mutant of Cdt1 (ND-Cdt1) with a removed PCNA interacting protein (PIP) degron that is required for PCNA binding and CRL4^Cdt2^-mediated degradation, and a mutated Cy motif that is required for its degradation by SCF^Skp2^ (Figure 2D and S2D) (Havens and Walter, 2009; Pozo et al., 2018; Sakaue-Sawano et al., 2017). Like full-length Cdt1, we find that ND-Cdt1 suppresses EdU incorporation (Figure 2E). Critically, inhibition of DNA synthesis did not prevent the firing of origins since CRL4^Cdt2^ was still activated similarly to control cells (Figure S2E). Furthermore, continued expression of ND-Cdt1 persistently inhibited EdU incorporation and prevented progression through S phase as measured by DNA content (Figures 2F, S2F and S2G). To ensure that ND-Cdt1 did not interfere with origin licensing, we measured chromatin-bound MCM2 as a measure of origin licensing (Håland et al., 2015; Matson et al., 2017) and found no change (Figure S2H). The inhibition of DNA replication by ND-Cdt1 was also observed in U2OS and HeLa cells, arguing that this inhibition is not cell-type specific and occurs in both non-transformed and transformed cells (Figures S2I and S2J).

As an additional control, we confirmed that endogenous Cdt1, not just overexpressed Cdt1, could inhibit DNA synthesis when it fails to be degraded in S phase. To prevent the degradation of Cdt1 in S phase, we acutely treated cells with MLN-4924, which blocks the activity of cullin-RING E3 ubiquitin ligases, including CRL4^Cdt2^ and SCF^Skp2^ (Figure S3A) (Lin et al., 2010). These cells had suppressed EdU incorporation following S phase entry (Figure 3A, siCtrl conditions). Similar to overexpressed Cdt1-mCherry, we found that MLN-4924 produced an increase in a population of cells with chromatin-bound PCNA and low EdU incorporation (Figure 3B, siCtrl conditions). Knockdown of Cdt1 partially rescued EdU incorporation, while knockdown of p21, another protein stabilized by MLN-4924 (Lan et al., 2016), did not, indicating that the suppression of DNA synthesis was mediated by the stabilized Cdt1 (Figure S3B). We conclude that both overexpressed and endogenous Cdt1 can suppress DNA synthesis during S phase. These findings provide a potential explanation of how cells avoid re-replication during the overlap period when Cdt1 is present together with fired origins in early S phase, as the amount of synthesized DNA, the substrate of re-replication, at these fired origins would be reduced until Cdt1 is fully degraded (Figure 3C).

**Figure 3.**
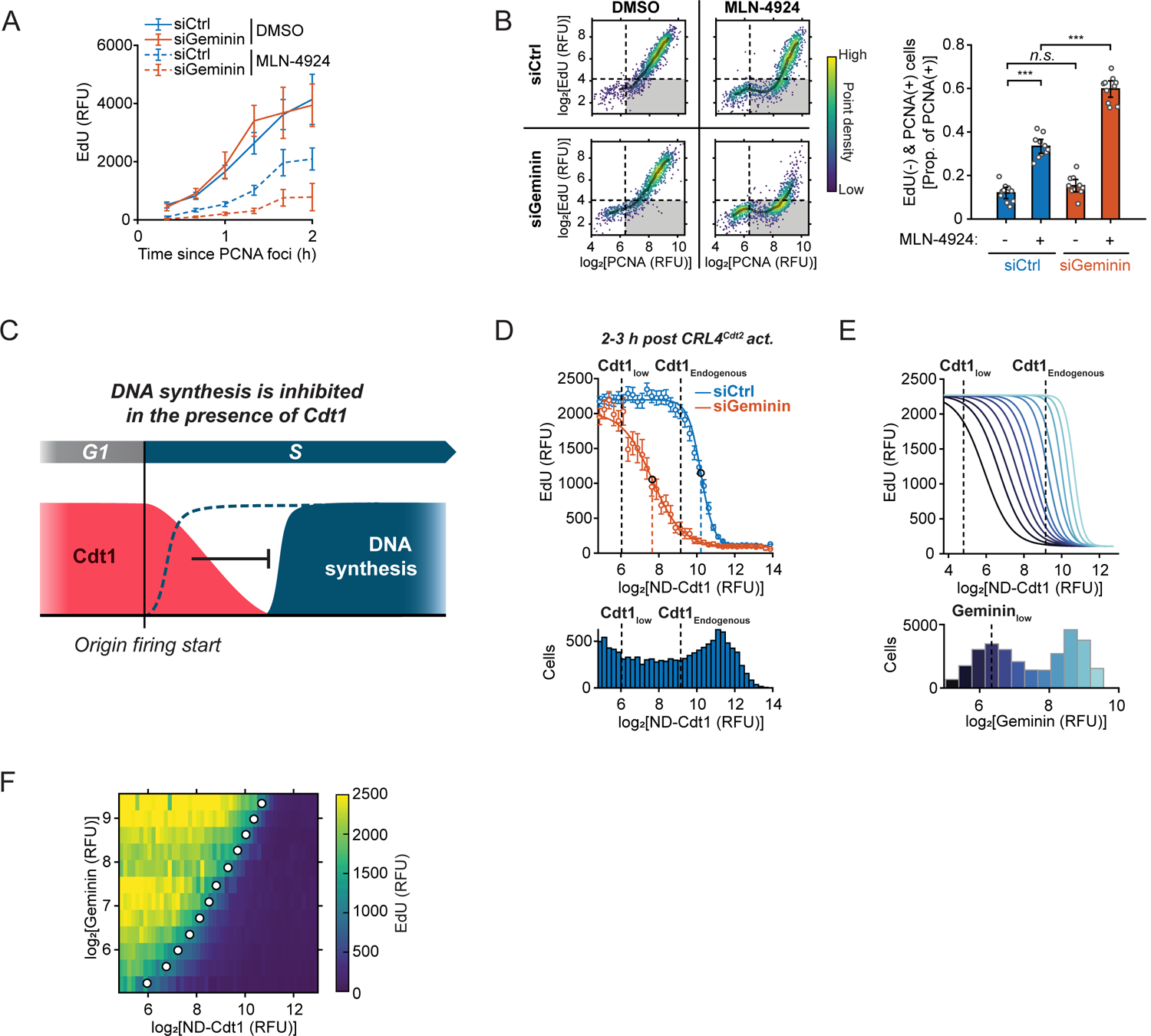
Endogenous Cdt1 can inhibit DNA synthesis and is counteracted by Geminin. **(A)** RT-QIBC of EdU incorporation, aligned to S phase start (PCNA foci) in cells treated with MLN-4924. Cells were transfected with siRNA for 4 h and then treated with 2 µM MLN-4924 for 3.5 h during live-cell imaging. Points and error bars are mean ± 2×SEM for each time point (n ≥8 for all timepoints, n ≥ 613 total for all conditions). **(B)** RT-QIBC of EdU incorporation and chromatin-bound PCNA in siRNA-treated, mitogen-released cells with the APC/C reporter and a Cyclin E/A-CDK activity reporter (see Figure S5A), fixed after 18 h. MLN-4924 was added 4 h prior to fixation. Cells 2-3 h after APC/C^Cdh1^ inactivation with Cyclin E/A-CDK activity ≥ 0.8 were analyzed. Cells were pooled from 10 replicate wells. Left: Dashed lines are thresholds based on G1 levels, and shaded curve is median in bins of PCNA levels (n ≥ 1,369 cells). Shaded quadrant contains PCNA positive and EdU negative cells. Right: Proportion of PCNA positive cells that are EdU negative. Points represent proportion in each of 10 wells. Error bars are mean ± 2×SEM. Two-sample *t*-test *p*-values: siCtrl DMSO vs. siCtrl MLN-4924 (7.8×10^-6^), siCtrl DMSO vs. siGeminin DMSO (.38), siCtrl MLN-4924 vs. siGeminin MLN-4924 (2.3×10^-5^). **(C)** DNA synthesis is inhibited in the presence of Cdt1 after origin firing. **(D-F)** RT-QIBC of EdU incorporation in siRNA-treated, mitogen-released cells with N-CRL4^Cdt2^ reporter. Cells were induced for ND-Cdt1 and fixed after 18 h. Representative of 3 independent experiments. **(D)** Top: Dose-response comparison of EdU incorporation. Points and error bars are mean ± 2×SEM for bins of ND-Cdt1 expression (bins ≥ 75 cells, 12,039 cells total siCtrl, 4,573 cells total siGeminin). Cells were stratified according to ND-Cdt1 expression levels, and a Hill equation was fit to the single-cell measurements. Maximum EdU inhibition was estimated from Hill equation fit to be 22.0-fold (siCtrl) and 23.0-fold (siGeminin). Dashed lines represent the IC_50_ (siCtrl: 10.2, siGeminin: 7.7) and Hill coefficients were 4.2 (siCtrl) and 1.8 (siGeminin). Bottom: Corresponding cell counts for bins of ND-Cdt1 levels in cells. Thresholds for equivalent ND-Cdt1 expression to endogenous Cdt1 (Cdt1_Endogenous_) and no ND-Cdt1 expression (Cdt1_low_) were calculated from Figure S2F. **(E)** Top: Fit ND-Cdt1 dose-response curves at 12 levels of Geminin expression. Bottom: Bins of Geminin expression selected for each of the 12 fits. Dashed line is threshold for low Geminin levels. See Figure S3C for siGeminin pooling to generate range of Geminin expression and Figure S3D for individual fits. n ≥ 660 cells per fit. **(F)** Heatmap of median EdU incorporation (color) at a given Geminin and ND-Cdt1 level, analyzed from same experiment as Figure 2E. Dots represent IC_50_ for each dose-response curve fit at each Geminin level. (n=29,350 cells total). See also Figure S3.

### Geminin counteracts the inhibition of DNA synthesis by Cdt1

When we inhibited Cdt1 degradation using MLN-4924, cells still started to increase EdU incorporation in S phase over time (Figure 3A). While Geminin is initially very low in early S phase, it gradually accumulates throughout S phase (Figure 1F). We considered whether Geminin binding to Cdt1 could ultimately abrogate the inhibitory action of Cdt1 on DNA synthesis as an additional mechanism by which cells could inactivate Cdt1 later in S phase. Consistent with this hypothesis, knockdown of Geminin suppressed EdU incorporation longer in cells treated with MLN-4924 (Figures 3A and 3B).

To better understand the role of Geminin, we determined how different levels of ND-Cdt1 regulate DNA synthesis in the presence or absence of Geminin. Cell lines with inducible ND-Cdt1 show variable expression between cells, and after computationally stratifying cells by their ND-Cdt1 expression, we found that EdU incorporation was inhibited in a dose-response manner taking the form of a Hill curve (Figure 3D). In cells 2-3 h post S phase entry, moderate ND-Cdt1 expression appeared to be buffered by Geminin and minimally inhibited EdU incorporation. In line with this, ND-Cdt1 suppressed DNA synthesis with a much lower IC_50_ with Geminin knocked down (Figure 3D).

To explore the relationship of Geminin to Cdt1-mediated inhibition of DNA synthesis further, we measured the dose-response of EdU to ND-Cdt1 at different Geminin levels, generated by treating cells with varying concentrations of Geminin siRNA (Figure S3C). We observed that DNA synthesis is progressively inhibited at lower ND-Cdt1 concentrations as Geminin decreases, with the IC_50_ decreasing linearly with Geminin levels (Figures 3E, S3D and S3E). This manifests as a ratiometric relationship between Cdt1 and Geminin, where double the ND-Cdt1 levels require double the Geminin levels to be neutralized (Figure 3F), consistent with Geminin’s role in physically binding to and inhibiting Cdt1 (Pozo and Cook, 2016).

Together, these analyses argue that Geminin not only inhibits Cdt1 licensing activity but also prevents Cdt1 from inhibiting DNA synthesis. This regulation allows DNA synthesis to proceed either after Cdt1 is degraded or after Geminin levels have sufficiently increased to inhibit Cdt1. Inhibition by Geminin likely becomes relevant in late S phase when Cdt1 is stabilized due to CRL4^Cdt2^ inactivation and Geminin levels are high, but DNA synthesis is still not complete (Pozo et al., 2018). However, during the overlap period of Cdt1 and fired origins in early S phase, there would not normally be enough Geminin to fully inhibit Cdt1, necessitating the suppression of DNA synthesis at fired origins by Cdt1 to deter re-replication.

### Cdt1 suppresses DNA synthesis during the overlap period of licensing and firing

If endogenous Cdt1 is indeed inhibiting DNA synthesis during early S phase, prematurely inactivating Cdt1 in G1 should accelerate the start of DNA synthesis. Since Cdt1 is required for origin licensing, we could not use long-term Cdt1 knockdown for these experiments. Instead, we made use of the licensing kinetics in MCF-10A cells, which complete the majority of origin licensing shortly after anaphase and then further boost licensing during G1 (Figures 4A and S4A). In this way, acutely inactivating Cdt1 during G1 would reduce but not prevent origin licensing, which cells are known to tolerate (McIntosh and Blow, 2012).

**Figure 4.**
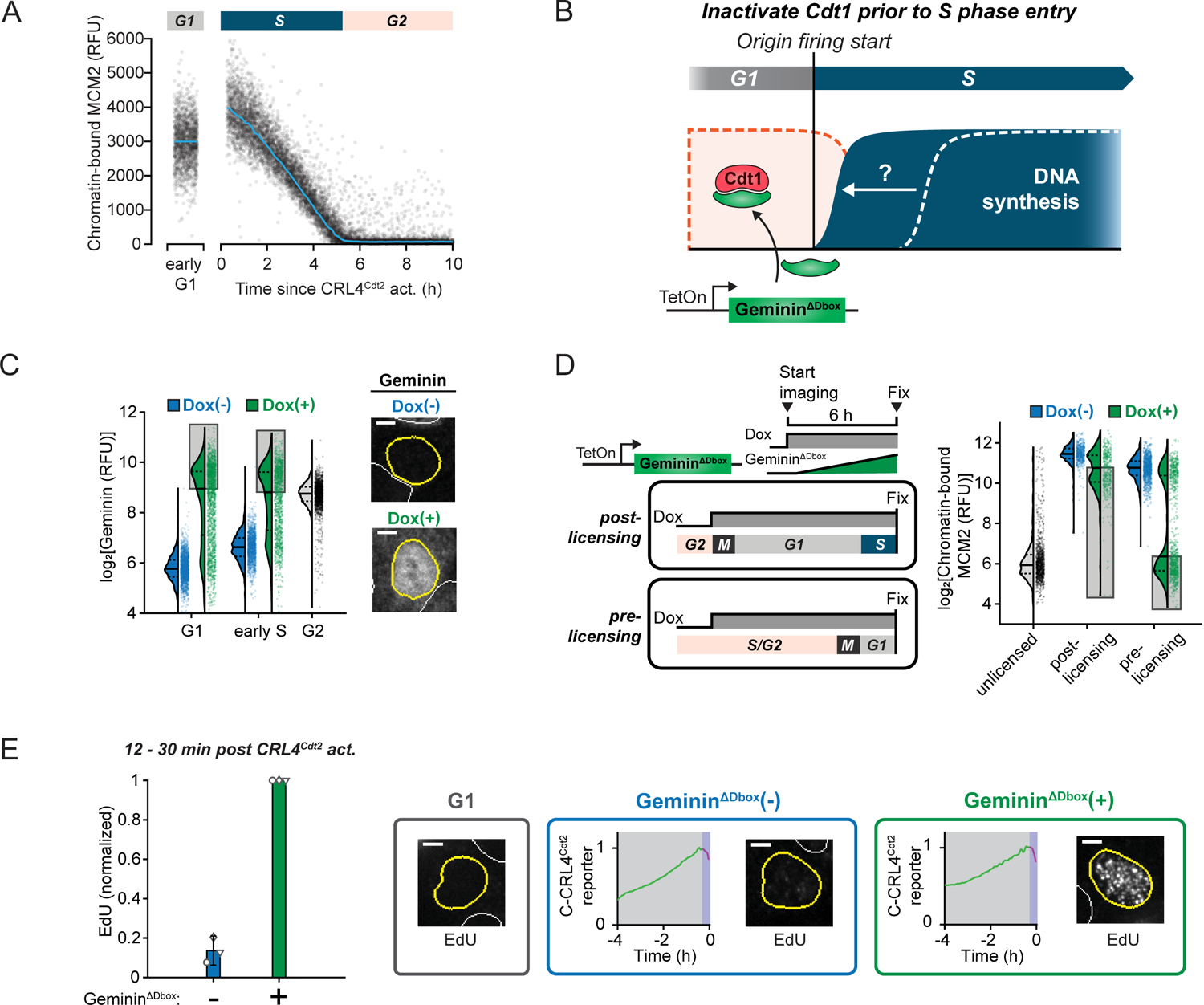
Cdt1 suppresses DNA synthesis during the overlap period of licensing and firing. **(A)** RT-QIBC of chromatin-bound MCM2, following S phase entry (C-CRL4^Cdt2^ reporter, n = 10,000 cells) and in early G1 (30 min – 1 h post anaphase, n = 1,500 cells). Pooled from 5 wells. Curves are median values. **(B)** Prematurely inactivating Cdt1 with doxycycline (Dox)-inducible Geminin^ΔDbox^ in G1 is predicted to accelerate the start of DNA synthesis after origin firing start. **(C-E)** Geminin^ΔDbox^ was induced with Dox during live-cell imaging of C-CRL4^Cdt2^ reporter for 6 h. **(C)** RT-QIBC of Geminin immunofluorescence (detects endogenous and Geminin^ΔDbox^) Left: G1 (1-2 h post anaphase), S phase (≤ 0.5 h in S phase) and G2 (4N DNA, EdU(-), no Dox). Shaded area is upper 50% of cells in Dox condition, which induce Geminin^ΔDbox^ to higher than normal G2 levels. Dashed and solid lines in violin plots are IQR and median, respectively. n ≥ 1,316 cells for each condition. Representative of 3 independent experiments. Right: Example Geminin signal in cells ∼1 h after anaphase in G1. 5 µm scale bar. **(D)** Left: Cells were identified in two groups of interest: 1) post-licensing addition in which Dox was added to cells ≤ 1 h before mitosis and which were in S phase for ≤ 1 h prior to fixation, and 2) pre-licensing addition in which Dox was added to cells at least 5 h before mitosis, blocking origin licensing. Right: RT-QIBC of chromatin-bound MCM2. Signal from unlicensed cells was estimated from G2 MCM2 signal (4N DNA and chromatin-bound PCNA negative). Shaded area is lower 50% of cells in Dox condition, corresponding to the approximately 50% of cells which induce Geminin^ΔDbox^ to higher than G2 levels (Figure 4C). Dashed and solid lines in violin plots are IQR and median, respectively. n ≥ 434 in all conditions. Representative of 2 independent experiments. **(E)** RT-QIBC of EdU incorporation in cells with Geminin^ΔDbox^ overexpressed, using the same gating as post-licensing cells from Figure 4D (Dox added ≤ 1h prior to mitosis). EdU was spiked onto cells 30 min prior to fixation during live-cell imaging. Cells that were in S phase for 12-30 min at fixation time, had not fully degraded Cdt1 were analyzed. Geminin^ΔDbox^(+) and Geminin^ΔDbox^(-) cells were selected based on Geminin stain. Left: For each of 3 independent experiments, the median of cells was taken (n ≥ 31 cells per replicate per condition; Geminin^ΔDbox^(-): 120 cells total; Geminin^ΔDbox^(+): 213 cells total) and normalized to the Geminin^ΔDbox^(+) condition. Error bars are mean ± 2×SEM (Geminin^ΔDbox^(-) cells are 13.6 ± 7.4% of Geminin^ΔDbox^(+) cells). See Figure S4E for estimated absolute DNA quantification. Right: representative EdU images and matching C-CRL4^Cdt2^ traces. Geminin^ΔDbox^(-) cell is 17.2 min into S phase, Geminin^ΔDbox^(+) cell is 16.9 min into S phase. 5 µm scale bar. See also Figure S4.

Since Geminin suppresses the Cdt1-mediated inhibition of DNA synthesis, we generated a cell line with Dox-inducible Geminin to prematurely inactivate Cdt1 during G1 (Figure 4B). To prevent Geminin degradation in G1 by APC/C^Cdh1^, we mutated the Geminin D-box motif (McGarry and Kirschner, 1998; Shreeram et al., 2002), and the resulting cell line induced Geminin^ΔDbox^ to levels higher than endogenous Geminin in G2 in ∼50% of G1 cells (Figure 4C).

When we acutely induced Geminin^ΔDbox^ in unsynchronized cells during live-cell imaging, we found that cells that divided shortly after Dox addition, and thus went through early G1 without Geminin^ΔDbox^, only had a moderate reduction in origin licensing at S phase entry, in line with the majority of origin licensing occurring shortly after anaphase in these cells (Figure 4D). In contrast, cells that received Dox well before mitosis and thus expressed Geminin^ΔDbox^ by the time they reached anaphase had completely inhibited origin licensing, indicating that Geminin^ΔDbox^ had fully inhibited Cdt1 in these cells. Therefore, we could examine cells where Geminin^ΔDbox^ was induced only during G1 (post-licensing Geminin^ΔDbox^ from Figure 4D) to determine when DNA synthesis starts if Cdt1 is inactivated before the start of S phase. In the first 30 min of S phase, cells with Cdt1 neutralized by Geminin^ΔDbox^ (Figure 4C, early S phase; Figures S4B and S4C) exhibited approximately 5 to 10-fold higher EdU incorporation than control cells (Figure 4E). We estimate that this increased EdU incorporation corresponds to approximately 12-18 megabases of DNA synthesized (Figure S4D and S4E). Thus, we conclude that Cdt1 limits the amount of synthesized DNA in early S phase, providing a protective mechanism against re-replication during an overlap period where Cdt1 is present together with fired origins.

### Cdt1 inhibits DNA synthesis independently of the intra-S phase checkpoint and re-replication

Next, we sought to determine the mechanism by which Cdt1 inhibits DNA synthesis. The intra-S phase checkpoint, which limits the rate of DNA synthesis and progression through S phase, can be activated in response to re-replication and DNA damage caused by Cdt1 dysregulation (Davidson et al., 2006; Liu et al., 2007; Truong and Wu, 2011; Vaziri et al., 2003). Alternatively, the addition of high levels of Cdt1 to replicating *Xenopus* egg extracts not only triggers checkpoint activation but also directly inhibits replication fork elongation (Nakazaki et al., 2016, 2017; Tsuyama et al., 2009), suggesting other plausible mechanisms by which Cdt1 could inhibit DNA synthesis.

We first focused on whether the intra-S phase checkpoint mediates the inhibition of DNA synthesis in early S phase by Cdt1. We overexpressed ND-Cdt1 in cells with Geminin knocked down to maximize the possibility of producing DNA damage and measured γH2AX, phospho-Chk1(S317) and phospho-Chk2(T68), markers of DNA damage and the intra-S phase checkpoint. Unexpectedly, we did not observe increases in these markers in response to ND-Cdt1 expression (Figure 5A).

**Figure 5.**
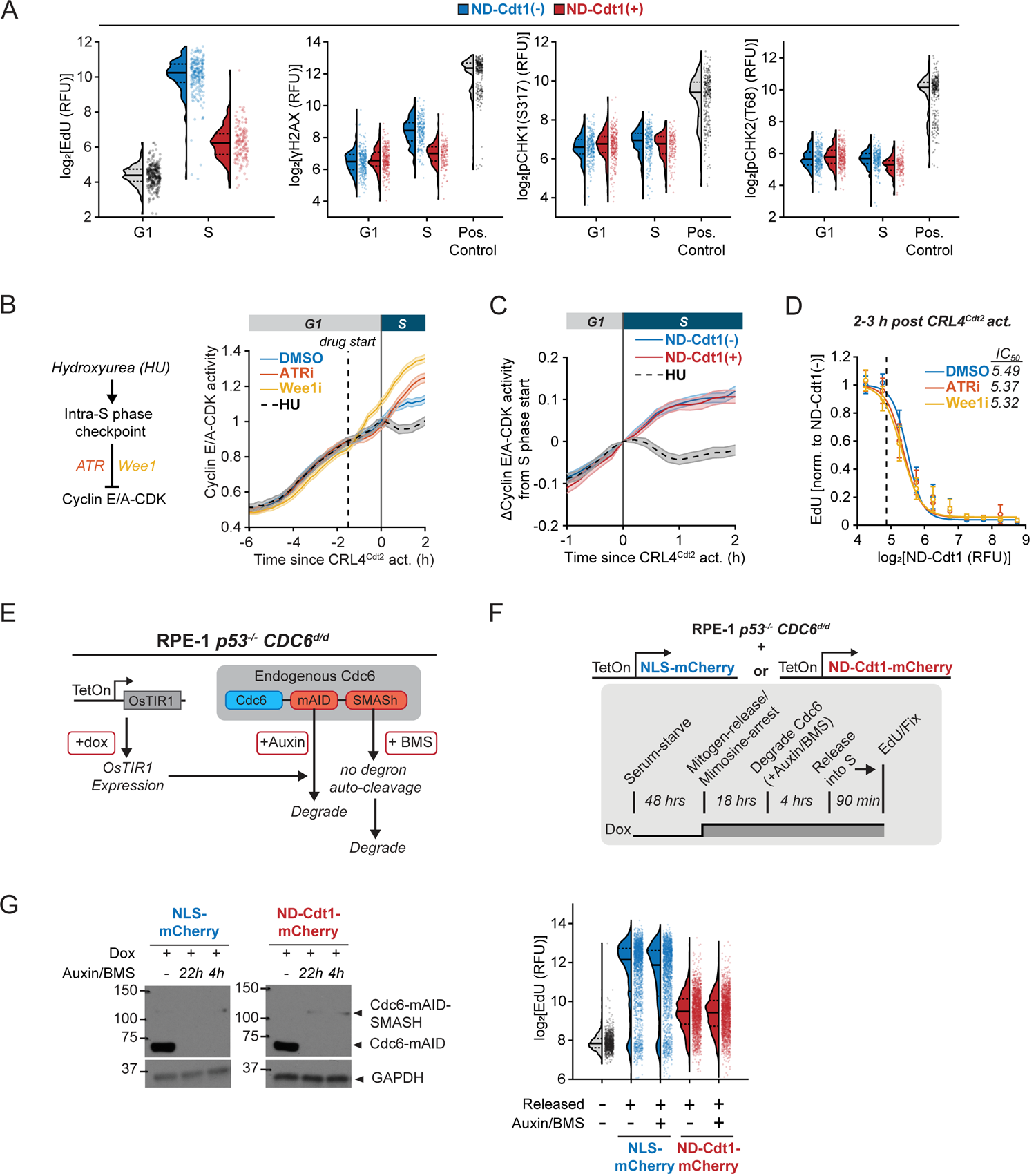
Cdt1 inhibits DNA synthesis independently of the global intra-S phase checkpoint and re-replication. **(A)** RT-QIBC of EdU incorporation, γH2AX, pChk1(S317) and pChk2(T68) in siGeminin-treated, mitogen-released cells with the N-CRL4^Cdt2^ reporter. Cells were treated with doxycycline (Dox) to induce ND-Cdt1 and fixed after 18 h. G1 cells had inactive CRL4^Cdt2^ with 2N DNA, while S phase cells were 1-2 h into S phase. Positive control for gH2AX, pChk1(S317) and pChk2(T68) staining was measured in cells treated with Wee1i for 4 h in S phase cells which caused an accumulation of DNA damage. n ≥ 146 cells for all conditions, representative of 2 independent experiments. **(B)** Left: Cyclin E/A-CDK activity is responsive to intra-S phase checkpoint activation. Right: Mitogen-released cells with N-CRL4^Cdt2^ and Cyclin E/A-CDK reporters were treated with Dox to induce ND-Cdt1 and imaged for 18 h. 14 h after release, 2 µM AZ-20 (ATRi), 1 µM MK-1775 (Wee1i) or 2 mM hydroxyurea (HU) were added to cells. Cells that received drug 1-2 h (dashed line is 1.5 h) prior to S phase entry were identified. Curves are mean traces, and the top and bottom of shaded area are 2×SEM. n = 167 (DMSO), 145 (ATRi), 239 (Wee1i) and 119 (HU) cell traces. Representative of 2 independent experiments. **(C)** Mitogen-released cells with N-CRL4^Cdt2^ and Cyclin E/A-CDK reporters were induced with Dox to express ND-Cdt1 and imaged for 18 h. 14 h after release 2 mM HU was added to control cells. Cells were stained for ND-Cdt1 at the end of the experiment, and ND-Cdt1(+) cells were selected for analysis. Change from Cyclin E/A-CDK activity at S phase start is shown. Lines and shaded area are mean ± 2×SEM. n ≥ 336 cell traces for all conditions. Representative of 2 independent experiments. **(D)** EdU incorporation dose-responses to ND-Cdt1 in the presence of 2 µM ATRi and 1 µM Wee1i. Mitogen-released, siGeminin treated cells with the N-CRL4^Cdt2^ reporter and ND-Cdt1 induced by Dox were fixed after 18 h. Drugs were added 4 h prior to fixation, and S phase cells that received the drug 1-2 h before S phase entry were selected. Points and error bars are mean ± 2×SEM for bins of ND-Cdt1 expression (bins ≥12 cells, and ≥770 cells total for all conditions). Estimated maximum EdU inhibition was 25.7-fold (DMSO), 17.6-fold (ATRi) and 17.0-fold (Wee1i). Representative of 2 independent experiments (same experiment as Figure 5A). **(E)** RPE-1 *p53^-/-^ CDC6^d/d^* cells had an auxin-inducible degron (mAID) and SMASh-tag knocked- in to both endogenous *CDC6* loci. Additionally, these cells contained a Dox-inducible OsTIR1 E3 ubiquitin ligase component which is required for mAID degradation. The addition of auxin to cells triggers the degradation of mAID containing proteins. The SMASh-tag contains degron which is auto-cleaved after protein translation by a protease domain. Addition of BMS-650032 (BMS) inhibits this auto-cleavage, resulting in protein degradation. Thus, addition of Dox/Auxin/BMS triggers a robust degradation of endogenous Cdc6. **(F)** Dox inducible constructs ND-Cdt1-mCherry or NLS-mCherry were introduced into RPE-1 *p53^-/-^ CDC6^d/d^* cells with the APC/C reporter. Cells were mitogen-released in the presence of mimosine and Dox for 18 h. Cdc6 was then degraded by adding auxin and BMS-650032 (BMS) for 4 h, and then cells were released from mimosine arrest for 1.5 h followed by an EdU pulse and fixation. **(G)** Left: Western blot of Cdc6 levels during experiment, comparing acute 4 h Cdc6 degradation to long-term Cdc6 degradation for 22 h from the time of serum release. Upper band is Cdc6 which has uncleaved SMASh-tag. Right: QIBC of EdU incorporation. S phase cells were selected based on having inactive APC/C^Cdh1^. Unreleased cells were not released from mimosine arrest. NLS-mCherry/ND-Cdt1-mCherry positive cells were chosen based on gates Figure S5D. n=1,120 cells (unreleased), 2,000 cells (other conditions). Representative of 2 independent experiments. See also Figure S5.

As an independent measure of checkpoint activation, we turned to a live-cell reporter of the activity of Cyclin E/A complexed with CDK2/1 (Cyclin E/A-CDK) (Figure S5A) (Chung et al., 2019; Spencer et al., 2013). Since Cyclin E/A-CDK activity is partially inhibited by the intra-S phase checkpoint, it can be used as a proxy for checkpoint activation (Daigh et al., 2018). Indeed, hydroxyurea reduces Cyclin E/A-CDK activity in S phase, while inhibitors of checkpoint mediators ATR or Wee1 increase Cyclin E/A-CDK activity (Figure 5B). However, ND-Cdt1 expression does not decrease Cyclin E/A-CDK activity, arguing against intra-S phase checkpoint activation (Figure 5C). As an additional test, we added ATR or Wee1 inhibitors to cells overexpressing ND-Cdt1 and found that ND-Cdt1 still suppressed EdU incorporation with the same IC_50_ (Figure 5D). Together, these results show that the intra-S phase checkpoint does not mediate the suppression of DNA synthesis by Cdt1.

It has also been suggested that re-replication can inhibit DNA synthesis independently of the intra-S phase checkpoint (Davidson et al., 2006; Neelsen et al., 2013). If re-replication is necessary for the suppression of DNA synthesis by Cdt1, blocking licensing activity, which is necessary for re-licensing and re-replication, should rescue DNA synthesis. To inhibit licensing, we used a previously developed RPE-1 *p53^-/-^* cell line with mAID and SMASh-tag inducible degrons knocked-in to both copies of *CDC6* (referred to here as *CDC6^d/d^*), another essential licensing factor (Figure 5E) (Lemmens et al., 2018). In this cell line, Cdc6 can be rapidly degraded to very low levels within 4 h (Figures 5F and 5G). Control experiments confirmed that degrading Cdc6 in cells entering the cell cycle from an unlicensed G0 inhibited origin licensing, resulting in a strong suppression of EdU incorporation (Figure S5B).

To determine whether re-licensing is required for Cdt1-mediated inhibition of DNA synthesis, we introduced Dox-inducible constructs for ND-Cdt1-mCherry and control NLS-mCherry into the *CDC6^d/d^* cell line. We synchronized cells in late G1 by releasing cells from G0 into a mimosine arrest, which blocks cells after origin licensing (Kubota et al., 2014). In the final 4 h before releasing cells from mimosine arrest into S phase, we could degrade Cdc6 to prevent further licensing during S phase and compare cells that expressed ND-Cdt1 or the control construct (Figures 5F, S5C and S5D). ND-Cdt1-mCherry inhibited EdU incorporation following mimosine release, regardless of Cdc6 degradation, arguing that re-licensing and re-replication are not necessary for Cdt1 to inhibit DNA synthesis (Figure 5G). Furthermore, this cell line does not have functional p53, which has also been implicated in the DNA damage response to re-replication (Vaziri et al., 2003). We conclude that re-replication, p53, and intra-S phase checkpoint activation are not required for the Cdt1-mediated inhibition of DNA synthesis in early S phase, which argues that Cdt1 directly suppresses DNA synthesis.

### Cdt1 inhibits replication fork elongation while permitting origin firing

We next determined whether Cdt1 suppresses DNA synthesis by inhibiting origin firing or by inhibiting replication fork elongation after origin firing, as has been suggested to occur in *Xenopus* egg extracts after the addition of excess Cdt1 (Nakazaki et al., 2016; Tsuyama et al., 2009). To quantify origin firing and recruitment of replication factors to the replication fork in the presence of ND-Cdt1, we measured chromatin-bound replication factors CDC45, TIMELESS, DNA polymerases epsilon, alpha and delta (Pol ε, α and δ), and PCNA (Figures 6A and S6B). Replication factors that are part of or bind to the CMG helicase (CDC45, TIMELESS, Pol ε and Pol α) did not have impaired chromatin association in the presence of ND-Cdt1, while Pol δ, which synthesizes lagging strands, and PCNA were present but approximately 50% reduced (Figures 6A, 6B and S6A-C) (Burgers and Kunkel, 2016). These findings are consistent with our observation that CRL4^Cdt2^, which depends on chromatin-bound PCNA, is still activated in the presence of ND-Cdt1 (Figure S2E).

**Figure 6.**
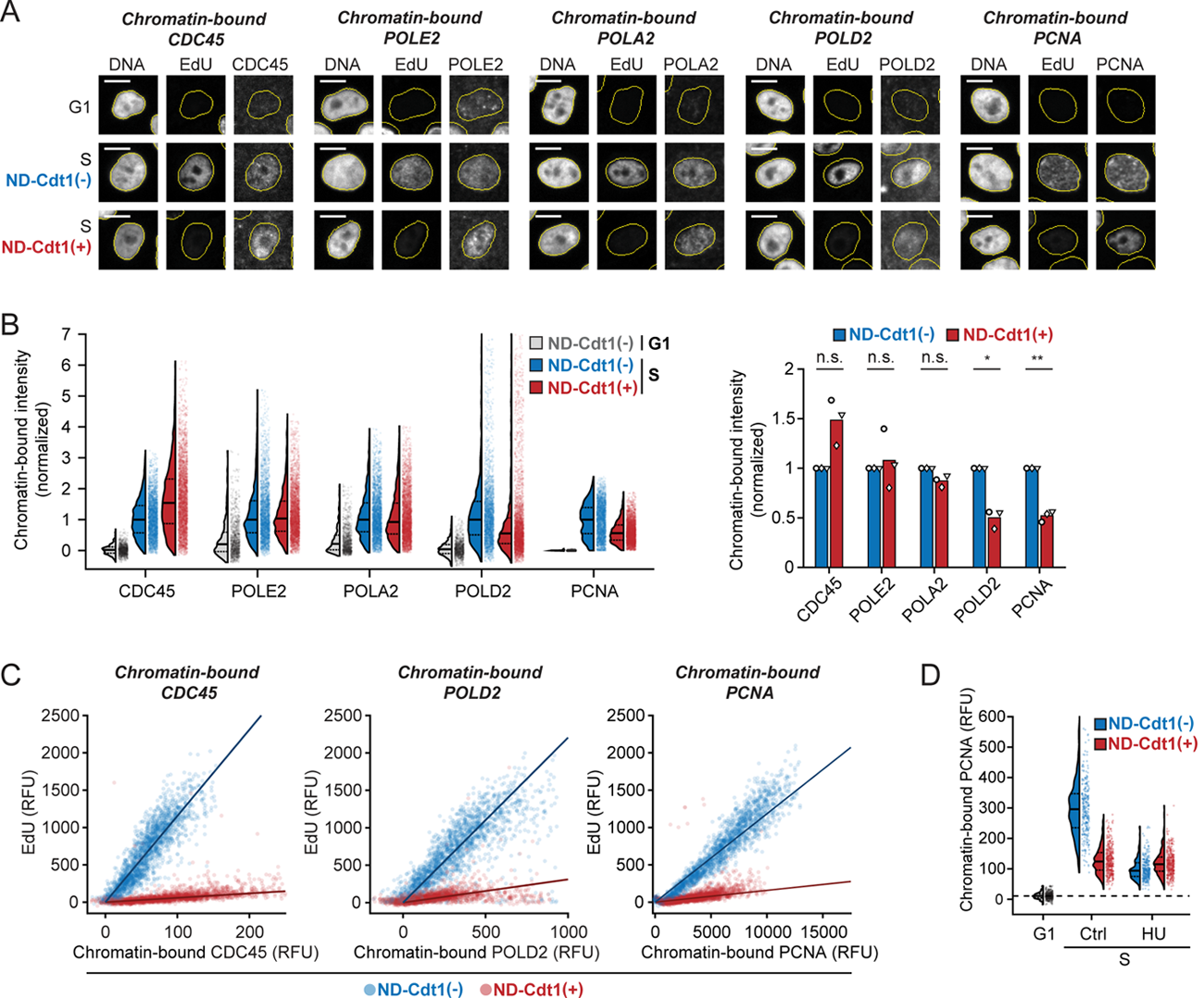
Cdt1 inhibits replication fork elongation while permitting origin firing. **(A-C)** RT-QIBC of chromatin-bound replisome components in cells with doxycycline (Dox)-inducible ND-Cdt1-mCherry. Mitogen-released cells with the APC/C reporter were induced with Dox and released for 18 h. G1 cells had active APC/C^Cdh1^ without chromatin-bound PCNA (co-stained with all proteins). S phase cells were 1-3 h post APC/C^Cdh1^ inactivation with chromatin-bound PCNA. Cells which expressed ND-Cdt1-mCherry during live imaging were selected for ND-Cdt1(+). Representative of 3 independent experiments. TIMELESS analysis in Figures S6B and S6C. **(A)** Representative cell images of EdU and chromatin-bound CDC45, Pol ε (POLE2), Pol α (POLA2), Pol δ (POLD2) and PCNA. Scale bar = 10 µm. **(B)** Quantification of chromatin-bound replisome components. G1 mode intensities were subtracted off signals and values were normalized to ND-Cdt1(-). Left: G1 (in absence of ND-Cdt1-mCherry) vs. S phase. Dashed and solid lines in violin plots are IQR and median respectively. n = 2,000 cells per condition. Right: Summary of median values from 3 independent experiments of left panel. One-sample Student’s t-test was performed on normalized Dox cells. *p*-values CDC45 (n.s.): 6.94×10^-2^, POLE2 (n.s.): .693, POLA2 (n.s.): 6.41×10^-2^, POLD2 (*): 1.21×10^-2^, PCNA (**): 4.3×10^-3^. **(C)** Analysis of EdU incorporation relative to chromatin-bound CDC45, POLD2 and PCNA. Fit line is from linear regression (n = 2,000 cells each condition). Other stains and summary in Figures S6D and S6E. **(D)** RT-QIBC of chromatin-bound PCNA. Mitogen-released cells with the APC/C reporter were induced with Dox and released for 18 h. During final 4 h of imaging, cells were treated with 2 mM hydroxyurea (HU) and then fixed. G1 cells had active APC/C^Cdh1^ without chromatin-bound PCNA, while S phase cells were fixed 2-3 h after APC/C^Cdh1^ inactivation and were chromatin-bound PCNA positive. n ≥ 281 cells for all conditions. Representative of 3 independent experiments. See also Figure S6.

To determine if the reduced levels of chromatin-bound Pol δ and PCNA we observe are responsible for the inhibition of DNA synthesis, we simultaneously analyzed the chromatin-bound replication factors together with EdU incorporation. In control conditions, there is a linear relationship between the level of each of the chromatin-bound proteins (CDC45, TIMELESS, Pol ε Pol α, Pol δ and PCNA) and EdU incorporation, indicating that each chromatin-bound protein signal is normally proportional to the number of active replication forks (Figures 6C and S6D). However, EdU incorporation is greatly reduced at matching levels of chromatin-bound protein in the presence of ND-Cdt1, suggesting that the same number of replication forks synthesize less DNA (Figures 6C and S6E). This is true even for Pol δ and PCNA, where EdU incorporation is much lower than would be expected given a 50% reduction in chromatin-bound levels. These results are consistent with Cdt1 inhibiting replication fork elongation at fired origins.

Since the lagging strand of the replication fork is bound by PCNA and Pol δ, which are eventually removed after the completion of Okazaki fragments (Burgers and Kunkel, 2016; Lee et al., 2013), we hypothesized that the reduced amounts of chromatin-bound PCNA and Pol δ we measure in the presence of ND-Cdt1 are a consequence of reduced fork elongation, rather than a reduced number of replication forks. In line with this, when we used hydroxyurea to block elongation of replication forks, we found lower chromatin-bound PCNA, which was not further lowered in cells also expressing ND-Cdt1 (Figure 6D). Such a reduction of PCNA at stalled forks has previously been reported (Sirbu et al., 2011; Yu et al., 2014). In aggregate, these results show that Cdt1 does not prevent the process of origin firing and formation of the replication fork, but rather inhibits replication fork elongation, which results in suppressed DNA synthesis.

### Cdt1 inhibits CMG helicase progression through its MCM-binding domains

Replication fork elongation primarily can be suppressed by inhibiting the replicative DNA polymerases or by inhibiting the progression of the CMG helicase, which is responsible for unwinding double-stranded DNA. Cdt1 contains a high-affinity interaction with PCNA through its PIP degron, and it has been suggested that PIP degron-containing proteins can interfere with the binding of polymerases to PCNA and thereby inhibit DNA synthesis (Mansilla et al., 2020; Tsanov et al., 2014). However, ND-Cdt1 has its PIP degron removed, arguing against polymerase inhibition. On the other hand, the addition of high levels of Cdt1 to replicating *Xenopus* egg extracts has been suggested to impair CMG helicase progression (Nakazaki et al., 2016, 2017). With these results in mind, we considered whether Cdt1 suppresses DNA synthesis in early S phase by inhibiting polymerases or CMG helicase progression.

A distinguishing feature of an inhibitor that blocks polymerases is that it triggers the accumulation of single-stranded DNA (ssDNA) at the replication fork, as CMG unwinds DNA that the polymerases are not able to fill (Figure 7A) (Nakazaki et al., 2016; Toledo et al., 2013; Zeman and Cimprich, 2014). Thus, we examined the chromatin-bound levels of ssDNA-binding protein RPA together with ND-Cdt1. During an unperturbed S phase, there is an increase in chromatin-bound RPA compared to basal G1 levels due to the normal production of ssDNA at replication forks (Figures 7B and S7A). However, cells with ND-Cdt1 have diminished RPA binding compared to control cells despite having similar, if not larger, amounts of origins fired, as measured by chromatin-bound CDC45. ND-Cdt1 also negates the increase in chromatin-bound RPA in response to hydroxyurea and ATR inhibitor co-treatment, which is known to generate a large increase in chromatin-bound RPA due to polymerase inhibition (Figures 7B and S7A) (Toledo et al., 2013). These results suggest that the CMG helicase unwinds less DNA in the presence of Cdt1. These findings are supported by a study in *Xenopus* egg extracts, where the addition of Cdt1 reduced the amount of chromatin-bound RPA (Nakazaki et al., 2016). We conclude that Cdt1 inhibits CMG helicase progression rather than polymerase activity in early S phase.

**Figure 7.**
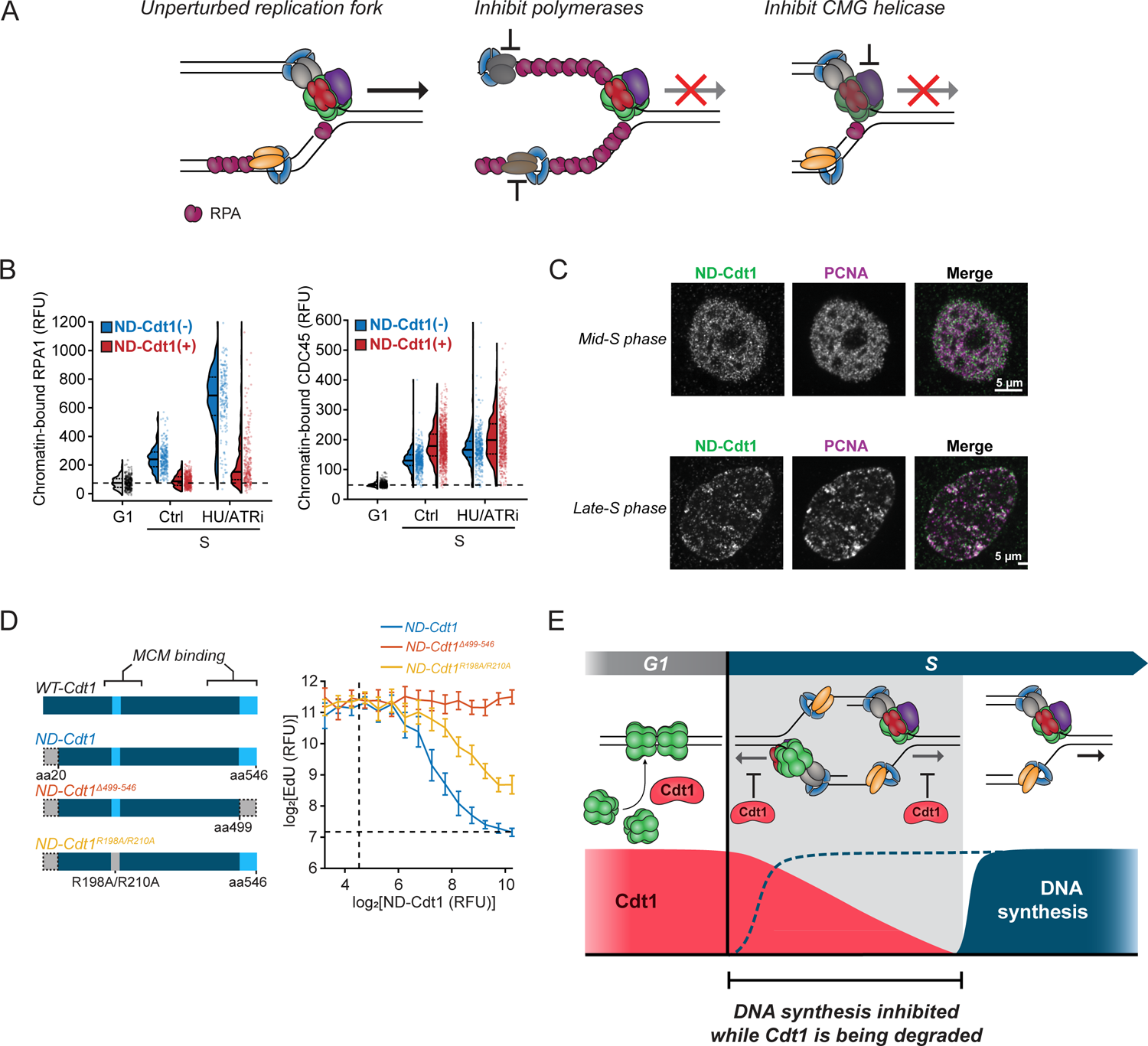
Cdt1 inhibits CMG helicase progression through its MCM-binding domain. **(A)** Replication fork progression can be inhibited by inhibiting CMG helicase progression or inhibiting DNA polymerases. **(B)** RT-QIBC of chromatin-bound RPA1 and CDC45. Mitogen-released cells with the APC/C reporter were released for 18 h and ND-Cdt1-mCherry was induced with doxycycline (Dox). During the final 4 h of imaging, cells were treated with 2 mM hydroxyurea (HU) and 2 µM AZ-20 (ATRi). G1 cells had active APC/C^Cdh1^ without chromatin-bound PCNA, while S phase cells were fixed 2-3 h after APC/C^Cdh1^ inactivation and were chromatin-bound PCNA positive. n ≥ 182 cells (RPA1) and n ≥ 504 cells (CDC45) for all conditions. Representative of 3 independent experiments. **(C)** Localization of chromatin-bound ND-Cdt1. Mitogen-released cells were treated with Dox and siGeminin to prevent the inactivation of ND-Cdt1 by Geminin. Cells were then pre-extracted after 18 h and co-stained for ND-Cdt1 (HA-tag) and PCNA. Cells were imaged using SoRa confocal microscopy and staged as either mid or late S phase based on the pattern of PCNA. Each image representative of ≥ 5 cells. **(D)** Left: MCM-binding region mutants of ND-Cdt1 were generated and introduced into cells in Dox-inducible constructs. WT-Cdt1 = wild-type Cdt1. Right: Mitogen-released cells with the N-CRL4^Cdt2^ reporter, treated with siGeminin and induced with Dox, were imaged for 18 h and then fixed. RT-QIBC of EdU incorporation and ND-Cdt1 staining (HA-tag) was performed and dose-responses of EdU 1-2 h after S phase entry were made. Points and error bars are mean ± 2×SEM in bins of ND-Cdt1 expression (n ≥26 cells for all bins n ≥ 1,048 cells total for each condition). Dashed lines were means calculated from cells uninduced with Dox (for ND-Cdt1) or G1 cells (for EdU). **(E)** Cdt1 degradation by CRL4^Cdt2^ is triggered by the start of origin firing. As Cdt1 is degraded to low levels, it inhibits CMG helicase progression to limit synthesized DNA production while Cdt1 is still present. DNA synthesis then commences in full following Cdt1 degradation. See also Figure S7.

As part of its role in origin licensing, Cdt1 directly binds to soluble MCM helicases through two MCM-binding regions, which results in a conformational change in the MCM helicases that allows them to be loaded onto origins (Frigola et al., 2017; Pozo and Cook, 2016). In *Xenopus* egg extracts, truncations in Cdt1 that overlapped with these regions interfered with the inhibition of DNA synthesis by Cdt1 (Nakazaki et al., 2017). We hypothesized that the same Cdt1-MCM binding interaction that occurs during licensing might also occur at the activated CMG complex, of which the MCM helicase is a component, and inhibit its progression. To test this hypothesis, we first examined the chromatin-binding of ND-Cdt1 using confocal microscopy. Strikingly, ND-Cdt1 colocalizes with replication foci (marked by chromatin-bound PCNA) during both early and late S phase (Figure 7C). Since ND-Cdt1 cannot bind PCNA due to its lack of PIP degron, we do not attribute the co-localization to direct binding to PCNA, but rather attribute it to CMG, which is expected to overlap with PCNA at the resolution limit of light microscopy.

We then tested whether the MCM-binding regions of Cdt1 are necessary for Cdt1 to inhibit DNA synthesis in human cells. The first MCM binding region is found at its C-terminus, while a second MCM interaction interface was identified near R210 of human Cdt1 (Pozo and Cook, 2016; Pozo et al., 2018). We overexpressed ND-Cdt1 with either a truncation at residue 498 in the C-terminal MCM-binding domain (ND-Cdt1^Δ499-546^), which abolishes licensing and MCM-binding (Teer and Dutta, 2008), or the point mutations R198A/R210A in the other interface (ND-Cdt1^R198A/R210A^), which severely diminishes licensing activity (Marco et al., 2009). We examined their inhibitory effect on DNA synthesis with Geminin knocked down to exclude differential Geminin regulation of the mutants and found that ND-Cdt1^Δ499-546^ cannot suppress EdU incorporation at all, while ND-Cdt1^R198A/R210A^ still inhibited EdU incorporation, albeit with an IC_50_ approximately double that of normal ND-Cdt1(Figure 7D). Our observation that Geminin, which prevents the binding of Cdt1 to MCM helicases (Pozo and Cook, 2016), also prevented Cdt1-mediated inhibition of DNA synthesis (Figure 3D) is consistent with this mutant analysis. Together, these experiments indicate that Cdt1 inhibits CMG helicase progression through the same MCM binding regions it uses for licensing.

## Discussion

Our study focused on the fundamental problem in eukaryotic DNA replication of duplicating the genome precisely once every cell cycle. It is generally thought that the solution to this problem is the strict temporal separation of origin licensing from origin firing to prevent re-replication. Vertebrate licensing regulation is centered around the inhibition of licensing factor Cdt1 from S phase entry to anaphase, through its inhibition by Geminin and Cyclin A and degradation by CRL4^Cdt2^ and SCF^Skp2^ (Pozo and Cook, 2016). However, only CRL4^Cdt2^ appears to act in early S phase, and given the dependence of CRL4^Cdt2^ activity on PCNA bound to replication forks, it has been noted this mechanism cannot fully separate licensing and firing in early S phase (Arias and Walter, 2007; Havens and Walter, 2011; Reusswig and Pfander, 2019). Considering the large number of replication origins, even a short overlap of Cdt1 and fired origins while Cdt1 is being degraded could allow for the re-licensing of DNA, leaving an open question of how cells might prevent re-replication during this vulnerable period.

In this work, we identified an overlap period of Cdt1 with fired origins in early S phase lasting approximately 30 min in human cells, during which Geminin and Cyclin A are still very low. Strikingly, we find that Cdt1 inhibits DNA synthesis during this overlap period, and this inhibition is only relieved once Cdt1 is fully degraded or Cdt1 becomes inhibited by increased expression of Geminin. Cdt1 suppresses CMG helicase progression, and thus replication fork elongation, at fired origins through its MCM-binding domains. By delaying replication fork elongation after origin firing, Cdt1 allows its own degradation by CRL4^Cdt2^ to initiate in response to origin firing while simultaneously reducing the production of synthesized DNA, which is the substrate of re-replication. Importantly, this mechanism is robust towards changes in Cdt1 expression levels, as cells with higher amounts of Cdt1 that take longer to degrade would suppress DNA synthesis longer. This protective mechanism could be particularly relevant in embryonic stem cells and cancer cells, which can have elevated Cdt1 levels (Matson et al., 2017; Truong and Wu, 2011).

Previous studies have identified responses to re-replication and DNA damage in human cells that reduce DNA synthesis in response to aberrant Cdt1 regulation (Truong and Wu, 2011). Critically, such mechanisms require re-replication to be produced before their activation and only minimize further damage, while CMG inhibition by Cdt1 can act before re-replication is produced. Cdt1 overexpression in human cells has been observed to impair S phase progression (Dorn et al., 2009; Takeda et al., 2005; Teer and Dutta, 2008), and we propose that Cdt1-mediated suppression of CMG progression can account for these observations in parallel with other mechanisms such as intra-S phase checkpoint activation. The finding that Cdt1 inhibits DNA synthesis raises the question of why dysregulation of Cdt1 has been previously shown to produce re-replication and DNA damage at all (Arias and Walter, 2005b, 2005a; Dorn et al., 2009; Klotz-Noack et al., 2012; Vaziri et al., 2003). A likely explanation is that the approximately 20-fold maximal inhibition of DNA synthesis by Cdt1 could still allow for enough residual DNA synthesis to produce re-replication over long periods. Furthermore, since overexpressed or dysregulated Cdt1 might be incompletely degraded, Cdt1 could be reduced to levels too low for effective suppression of DNA synthesis, but high enough for some re-licensing to occur over time. In support of this, non-degradable mutants of Cdt1 paradoxically produce less re-replication than wild-type Cdt1 (Takeda et al., 2005; Teer and Dutta, 2008).

Overall, a combined single-cell analysis of live- and fixed-cell microscopy enabled us to observe the dynamics of licensing regulation and DNA synthesis within the transition period in early S phase. Our study suggests a revision of the concept that origin licensing must strictly be separated from origin firing to avoid re-replication, and argues that human cells instead separate origin licensing from DNA synthesis in early S phase. Importantly, we identify that this separation is enforced by Cdt1 inhibiting the CMG helicase after origin firing. Previously identified re-replication prevention mechanisms center around the inhibition of licensing factors as cells enter S phase (Arias and Walter, 2007). In contrast, we identified a new class of licensing regulation whereby a licensing factor itself inhibits S phase progression. We propose that both classes of regulation are needed to safeguard genome integrity in early S phase.

## Acknowledgments

We thank Arne Lindqvist for providing RPE-1 *p53^-/-^ CDC6^d/d^* cells; Hana Sedlackova and Jiri Lukas for providing U2OS *CDC45-GFP* cells; Yilin Fan, Lindsey Pack, Anjali Bisaria, Damien Garbett, Steven Cappell, and Arnold Hayer for technical support; Meredith Weglarz and the Stanford Shared FACS Facility for cell sorting; Lindsey Pack, Yilin Fan, and Katherine Ferrick for critical reading of the manuscript; Karlene Cimprich, James Ferrell, Gheorghe Chistol, Daniel Jarosz and all members of the Meyer laboratory for helpful discussions. This work was funded by a National Institute of General Medical Sciences (NIGMS) R35 grant (5R35GM127026-05). N. R. was supported by an NSF Graduate Research Fellowship (DGE-1147470).

## Author Contributions

Conceptualization, N.R., M.C., and T.M; Methodology, N.R., M.C., and T.M; Software, N.R.; Formal Analysis, N.R.; Investigation, N.R., and M.C.; Data Curation, N.R.; Writing – Original Draft, N.R., and T.M.; Writing – Review & Editing, N.R., M.C., and T.M; Visualization, N.R.; Supervision, T.M; Funding Acquisition, T.M.

## STAR Methods

### Resource availability

#### Lead contact

Further information and requests for resources and reagents should be directed to and will be fulfilled by the lead contact, Tobias Meyer

## Materials availability

Plasmids and cell lines generated in this study are available upon request to lead contact.

## Data and code availability

Custom MATLAB image-processing pipeline and scripts used to generate the figures from this study have been deposited at Github and Zenodo (https://github.com/MeyerLab/image-analysis-ratnayeke-2021, https://doi.org/10.5281/zenodo.5037903<u>). Data from processed images to use with the code repository are available at Dryad (</u>https://doi.org/10.5061/dryad.4xgxd2599<u>). Raw imaging data acquired during this study have not been deposited in a public repository due to storage limitations but are available from the lead contact upon request.</u>

### Experimental model and subject details

#### Cell culture

All experiments were performed with MCF-10A human mammary epithelial cells (ATCC Cat# CRL-10317, RRID:CVCL_0598) unless otherwise noted. MCF-10A cells were cultured in DMEM/F12 growth media with HEPES (Gibco Cat# 11039047), supplemented with 5% horse serum (Gibco Cat# 16050122), 20 ng/mL EGF (PeproTech Cat# AF-100-15), 0.5 µg/mL hydrocortisone (Sigma: H0888), 100 ng/mL cholera toxin (Sigma Cat# C8052) and 10 µg/mL insulin (Sigma Cat# I1882). Cells were passaged using trypsin-EDTA (0.05%, Gibco Cat# 25300054) and trypsin was neutralized in DMEM/F12 supplemented with 20% horse serum. RPE-1 *p53^-/-^* cells with double-degron endogenous-tagged Cdc6 (RPE-1 *p53^-/-^ CDC6^d/d^*) were a kind gift from Arne Lindqvist (Lemmens et al., 2018) and cultured in DMEM/F12 with HEPES supplemented with 10% FBS (Sigma Cat# TMS-013-B). U2OS cells had endogenously GFP-tagged CDC45 (a kind gift from Jiri Lukas [Sedlackova et al., 2020]) and were cultured in DMEM growth media (Gibco Cat# 11995065) with 10% FBS. HeLa cells (ATCC Cat#CCL-20.2, RRID:CVCL_2260) were cultured in DMEM growth media with 10% FBS. For MCF-10A serum starvation, cells were cultured in starvation media (growth media without horse serum, EGF and insulin and supplemented with 0.3% BSA) after two washes of starvation media. For mitogen-release, starvation media was exchanged with growth media. All cells were cultured at 37°C and 5% CO2. For microscopy experiments, 96-well glass-bottomed plates (Cellvis Cat# P96-1.5H-N) were collagen-coated (Advanced Biomatrix Cat# 5005-B, 60 µg/mL dilution for at least 2 h) and cells were seeded into wells at least the night before performing experiments.

#### Cell line generation

Cell cycle reporter cell lines were generated using third-generation lentiviral transduction (Dull et al., 1998; Stewart et al., 2003). In short, lentivirus was produced in HEK-293T cells co-transfected with packaging plasmids pMDLg/pRRE (Addgene # 12251, RRID:Addgene_12251), pRSV-rev(Addgene # 1225, RRID:Addgene_12253), and pCMV-VSV-G (Addgene # 8454, RRID:Addgene_8454) together with the lentiviral plasmid with Lipofectamine 2000 (Thermo Cat# 11668019). 72 h after transfection, virus was collected from the supernatant, filtered with a .22 µm filter (Millipore Cat# SCGP00525) and concentrated using 100 kDa centrifugal filters (Millipore Cat# UFC910024). Virus was then transduced into cells in growth media. For constitutively expressed fluorescent constructs, positive fluorescent cells were sorted using a BD Influx cell sorter (performed in Stanford Shared FACS Facility), while Dox-inducible constructs (TetOn in pCW backbone with puromycin selection marker) were selected with 1 µg/mL puromycin until control cells died unless otherwise stated. TetOn cells were grown in the absence of Dox until the time of experiment unless otherwise stated.

All MCF-10A reporter cell lines were generated from a base cell line transduced with CSII-pEF1a-H2B-mTurquoise as a nuclear tracking marker. Cells with EYFP-PCNA or the APC/C reporter were generated by transducing cells with pLV-EYFP-PCNA or CSII-pEF1a-mVenus-hGeminin(1-110) respectively. Cells containing the APC/C reporter together with either N- or C-CRL4^Cdt2^ reporter were generated by transducing cells with bicistronic vector tFucci(CA)2/pCSII-EF (Sakaue-Sawano et al., 2017) or pLV-hCdt1(1-100)ΔCy-mCherry-P2A-mVenus-hGeminin(1-110) respectively. Cells with the Cyclin E/A-CDK reporter together with N-CRL4^Cdt2^ were created by transduction with CSII-pEF1a-hDHB(994-1087)-mVenus and pLV-mCherry-hCdt1(1-100)ΔCy.

pCW constructs (TetOn Dox-inducible) expressing HA or mCherry-tagged Cdt1 mutants or Geminin^ΔDbox^ were introduced into these fluorescent reporter cell lines in combinations found in the Table S1. For cells with the APC/C reporter and TetOn-Cdt1-mCherry, cells were transduced with CSII-pEF1a-mVenus-hGeminin(1-110), followed by pLV-rtTA3-IRES-Puro, and then pLV-TetOn-Cdt1-mCherry. Cell lines transduced with Cdt1-mCherry constructs were induced with 500 ng/mL Dox while serum starved to sort for expressing cells. mCherry positive cells were sorted, and media was then switched to growth media without Dox. MCF-10A cells containing a CDK4/6 reporter (not analyzed), Cyclin E/A-CDK reporter and APC/C reporter used in Figures S3A and S3B were described previously (Yang et al., 2020). RPE-1 *p53^-/-^ CDC6^d/d^* cells were transduced with with CSII-pEF1a-H2B-mTurquoise and CSII-pEF1a-mVenus-hGeminin(1-110), and then pLV-TetOn-ND-Cdt1-mCherry or pLV-TetOn-NLS-mCherry. U2OS *CDC45-GFP* cells were transduced with pCW-ND-Cdt1-mCherry-Puro. HeLa cells were transduced with CSII-pEF1a-H2B-mTurquoise and pLV-mCherry-hGeminin(1-110)-IRES-Puro.

### Method Details

#### Cell cycle reporters

Cell cycle reporters of CRL4^Cdt2^ and APC/C activity were used in this study. These reporters were originally developed as the two components of the FUCCI(CA) reporter system (Sakaue-Sawano et al., 2017). The CRL4^Cdt2^ activity reporter is based on a fragment of human Cdt1 corresponding to the amino acid 1-100, and this fragment is inactive with respect to origin licensing. The fragment Cdt1(1-100)^ΔCy^ contains a PCNA-interacting protein (PIP) degron, which mediates Cdt1 degradation by CRL4^Cdt2^ in response to PCNA at fired origins, and has a removed Cy motif to prevent degradation by SCF^Skp2^ (Sakaue-Sawano et al., 2017). The CRL4^Cdt2^ reporter is rapidly degraded to low levels at S phase start and reaccumulates at the start of G2. Conversely, the APC/C reporter is based on amino acids 1-110 of human Geminin fused to a fluorescent protein (either mVenus or mCherry), is degraded at anaphase by APC/C^Cdc20^ and then by APC/C^Cdh1^ throughout G1, and reaccumulates at the time of APC/C^Cdh1^ inactivation at the G1/S transition. APC/C^Cdh1^ inactivation represents a commitment point in the cell cycle and typically occurs near the time of S phase entry and DNA replication, though CRL4^Cdt2^ activation in response to origin firing is an explicit measure of S phase entry (Grant et al., 2018; Sakaue-Sawano et al., 2017). While these reporters are typically quantified by their presence or absence in single-timepoint measurements, when reporter fluorescence kinetics are measured in single cells using time-lapse microscopy, the precise time of CRL4^Cdt2^ activation (the start of degradation of the CRL4^Cdt2^ reporter) and APC/C^Cdh1^ inactivation (the stabilization of the APC/C reporter) can be identified.

In this study, we used two versions of the CRL4^Cdt2^ reporter, one of which is an N-terminal mCherry-tagged Cdt1(1-100)^ΔCy^ (referred to as the N-CRL4^Cdt2^ reporter), which is identical to the construct used in the FUCCI(CA) reporter system. Since the N-CRL4^Cdt2^ reporter is fused to a fluorescent protein on the N-terminus, the PIP degron is in the middle of the construct. Since Cdt1 naturally has an N-terminal PIP degron, we hypothesized that reversing the order of the fluorescent protein fusion in a reporter could confer a faster response to the initial origins that are fired in early S phase. As a result, we created a C-terminally tagged Cdt1(1-100)^ΔCy^ (referred to as the C-CRL4^Cdt2^ reporter). We found that C-CRL4^Cdt2^ responds with slightly faster kinetics at S phase start than N-CRL4^Cdt2^, which was necessary for looking within the first 15-30 min of S phase by RT-QIBC in Figures 1 and 4. However, both reporters are well suited for RT-QIBC looking at times after the first 15-30 min of S phase, and we use both reporters in this study. C-CRL4^Cdt2^ has a similar orientation to the PIP-FUCCI cell cycle reporter (Grant et al., 2018), which is based on Cdt1(1-17) and is also degraded throughout S phase. We consider the N-CRL4^Cdt2^ (originally used in the FUCCI(CA)) system, C-CRL4^Cdt2^ and PIP-FUCCI reporters to all be reporters of CRL4^Cdt2^ activity, and should all be suitable for use with RT-QIBC.

The cyclin E/A-CDK reporter is a translocation-based reporter which is phosphorylated by cyclin E or A complexed with CDK2 or CDK1 (referred to collectively as cyclin E/A-CDK) (Spencer et al., 2013). It is based on a fragment of human DNA helicase B (amino acids 994-1087), which is phosphorylated by cyclin E/A-CDK. When unphosphorylated in G0 and early G1, this reporter is localized in the nucleus, and as cyclin E/A-CDK activity increases throughout the cell cycle, the reporter becomes progressively localized to the cytoplasm due to increased phosphorylation. Thus, the cytoplasm to nuclear ratio of intensity is a readout of cyclin E/A-CDK activity.

### Plasmid generation

Plasmids generated in this study were assembled using Gibson assembly of PCR amplified inserts and restriction enzyme digested plasmid backbones. Human full-length Cdt1 was amplified out of MCF-10A cDNA for Cdt1 overexpression, and mutations and tags were introduced through primers or gene synthesis (IDT). ND-Cdt1 constructs were created through a truncation of wild-type Cdt1 (aa20-546) which removes the Cdt1 PIP degron. The Cdt1 Cy motif (aa68-70 of full-length Cdt1) was mutated to alanine (ΔCy) to prevent degradation by SCF^Skp2^. This sequence was fused at the C-terminus to a flexible linker, SV40 NLS and either an mCherry or HA tag. The Geminin^ΔDbox^ (human Geminin with R23A and L26A mutations) sequence was generated using gene synthesis and HA-tagged. For Dox-inducible TetOn constructs, PCR products were inserted into pCW backbone (derived from pCW-Cas9, a gift from Eric Lander & David Sabatini, Addgene plasmid # 50661, RRID:Addgene_50661), a bicistronic vector with a TetOn promoter driving gene expression in addition to a constitutive PGK promoter driven PuroR-T2A-rtTA. pC1-ND-Cdt1-mCitrine was created by cloning ND-Cdt1 into the pC1 backbone, derived from C1-F-tractin-mCitrine, (Bisaria et al., 2020). pLV-hCdt1(1-100)ΔCy-mCherry-P2A-mVenus-hGeminin(1-110) was generated from full-length Cdt1 and Geminin (Human ORFeome V5.1). N-CRL4^Cdt2^ reporter was amplified from tFucci(CA)2/pCSII-EF, and inserted into the pLV backbone to generate pLV-mCherry-hCdt1(1-100)ΔCy. pLV, CSII and pCW are lentiviral expression plasmids, while pC1 is a mammalian expression plasmid.

### siRNA and plasmid transfection

MCF-10A cells were transfected with siRNA using DharmaFECT 1 (Dharmacon Cat# T-2001-03) according to the manufacturer’s protocol using 20 nM siRNA and 1:500 diluted DharmaFECT 1 final concentration unless otherwise stated. Cells were incubated in transfection mixture for at 6-24 h in either growth or serum starvation media, followed by a media change. Pools of 3-4 siRNA oligos (ON-TARGETplus, Dharmacon) were used for siCtrl, siCdt1 and siGeminin. For siCdt1 and siGeminin, oligos that do not target hCdt1(1-100) and hGeminin(1-110) were selected to avoid knockdown of the CRL4^Cdt2^ and APC/C reporters, respectively. A list of siRNA oligos is in Table S1. HeLa cells were transiently transfected with pC1-ND-Cdt1-mCitrine plasmid using Lipofectamine 2000 according to the manufacturer’s protocol using 2 ng/µL final concentration of plasmid complexed with 1:400 diluted Lipofectamine 2000 final concentration. Media was exchanged with growth media after 2 h, and cells were then immediately live-imaged. siRNA and plasmids were both complexed in Opti-MEM serum-free media (Gibco Cat# 31985070).

### Drugs

Stock solutions of drugs were dissolved in DMSO (Sigma-Aldrich Cat# D2650) and used at the given working concentration unless otherwise stated: 2 mM hydroxyurea (HU, dissolved in water, Cayman Chemical Cat# 23725), 2 µM AZ-20 (ATRi, Cayman Chemical Cat# 17589), 1 µM MK-1775 (Wee1i, Cayman Chemical Cat# 21266), 1 µg/mL Doxycycline hyclate (Sigma-Aldrich Cat# D9891), 2 µM MLN-4924 (Abcam Cat# ab216470), 500 µM indole-3 acetic acid (auxin, MP Biomedicals Cat# 0210203705), 100 nM BMS-650032 (Adooq Bioscience Cat# A112955), 500 µM L-mimosine (20x stock solution dissolved in DMEM/F12, Cayman Chemical Cat# 14337). For release from mimosine arrest, cells were washed three times in growth media. For all experiments where drugs or Doxycycline were added to cells, DMSO (vehicle) was added to control cells, with the exception of HU, which was dissolved in water.

### Western blot

Cells were grown in 6-well plates. At the time of lysis, cells were washed in ice-cold PBS, lysed in 2x Laemmli sample buffer with 100 mM DTT and a cell scraper, passed through a 25G needle 10 times, and heated at 90°C for 5 min. Samples were then separated with SDS-PAGE using 7.5% Mini-PROTEAN TGX gels (Bio-Rad Cat# 4561025) in Tris/Glycine/SDS running buffer (Bio-Rad Cat#161-0772), followed by semi-dry transfer (Bio-Rad Trans-Blot SD, Cat# 1703940) onto 0.45 µm PVDF membranes (Millipore Cat# IPFL00010) with Tris/Glycine buffer (Bio-Rad Cat# 1610734) + 10% MeOH. Membranes were washed in TBST (20 mM Tris, pH 7.5, 150 mM NaCl, 0.1% Tween 20), blocked for 30 min in 5% milk in TBST, and incubated overnight with mouse anti-CDC6 antibody (1:500, Santa Cruz Biotech. Cat# sc-9964, RRID:AB_627236) or rabbit anti-GAPDH (1:1000, CST Cat# 5174, RRID:AB_10622025) in 5% BSA + 0.01% NaN_3_ in TBST. Membranes were then incubated in HRP secondary antibodies (1:5000, CST Cat# 7074, RRID:AB_2099233 or CST Cat# 7076, RRID:AB_330924) for 30 min, treated with chemiluminescent substrate (Thermo Cat # 34580) and detected on film (Thomas Sci. Cat# EK-5130).

### Fixed-cell sample preparation

#### General Protocol

Staining and imaging were performed in 96-well glass-bottomed plates (Cellvis Cat# P96-1.5H-N). Cells were fixed in 4% paraformaldehyde in PBS (diluted from Fisher Cat# NC1537886) for 10 min at room temperature followed by PBS wash. If cells expressed fluorescent proteins which spectrally overlapped with the fluorophores used in later steps, the fluorescent proteins were chemically bleached (Lin et al., 2015) in 3% H_2_O_2_ + 20 nM HCl in PBS for 1 h, washed in PBS, and checked under a microscope to ensure there was negligible residual signal. If fluorescent proteins needed to be quantified in fixed cells prior to immunofluorescence, cells were initially imaged before bleaching and reimaged after staining. For PCNA and CDC45 staining, cells were incubated in ice-cold methanol for 15 min after fixation and then washed in PBS. Cells were permeabilized in 0.2% Triton X-100 in PBS for 10 min and then blocked in blocking buffer A (10% FBS, 1% BSA, 0.1% Triton X-100, 0.01% NaN_3_ in PBS) for 1 h. Cells were then incubated with primary antibodies overnight in blocking buffer A at 4°C, washed in PBS, and then incubated with secondary antibodies in blocking buffer A for 1 h at RT. Cells were washed with PBS and then incubated in 1 µg/mL Hoechst 33342 (Invitrogen Cat# H3570) in PBS for 10 min, followed by a final PBS wash prior to imaging. Unless otherwise stated, all washes were done with an automated plate washer (aspirate to 50 µL, dispense 250 µL, repeated 9 times, BioTek 405 LS), or by hand (for pre-extracted samples, 3 washes aspirating all liquid).

#### Iterative immunofluorescence

If simultaneously staining for targets with antibodies of the same species, the iterative indirect immunofluorescence imaging (4i) technique was used to sequentially image multiple antibodies (Gut et al., 2018). In short, the first round of imaging was identical to the general immunofluorescence protocol, with the exception that cells after the post-Hoechst PBS wash were washed in ddH_2_O and then placed in imaging buffer (700 mM N-acetyl cysteine in ddH_2_O, pH 7.4, Sigma-Aldrich A7250). Cells were imaged and then washed in ddH_2_O. The prior-round antibodies were eluted by 3×10-min incubations in elution buffer, which consists of 0.5M glycine (Sigma-Aldrich), 3M urea (Sigma-Aldrich), 3M guanidinium chloride (Sigma-Aldrich) and 70 mM TCEP-HCl (Goldbio Cat#TCEP50) in ddH_2_O, pH 2.5, followed by a PBS wash. Cells were then checked under a fluorescence microscope to ensure proper elution. Cells were then blocked with blocking buffer B, consisting of 1% BSA in PBS supplemented with 150 mM maleimide (dissolved just prior to use, Sigma-Aldrich Cat# 129585) for 1 h and then washed in PBS. Cells were then blocked with blocking buffer A for 30 min, followed by primary antibody incubation, and the subsequent steps the same as in the first round, repeated as needed. Control wells leaving out primary antibodies were always included to ensure there was no residual signal from prior rounds of imaging.

#### Pre-extraction for chromatin-bound protein

If chromatin-bound proteins were being stained for, soluble proteins were extracted from cells. Just prior to fixation, media was aspirated off of cells, and the plate was placed on ice. Cells were incubated in ice-cold pre-extraction buffer, consisting of 0.2% Triton X-100 (Sigma-Aldrich Cat# X100) + 1x Halt Protease Inhibitor Cocktail (Thermo Cat# 78439) in selected aqueous buffer. For all proteins pre-extracted for except RPA1, pre-extraction buffer was made with PBS, while CSK buffer was used for RPA1, consisting of 10 mM PIPES (Sigma-Aldrich), 100 mM NaCl (Sigma-Aldrich), 300 mM sucrose (Sigma-Aldrich), 3 mM MgCl_2_ (Sigma-Aldrich) at pH 7.0. After a set extraction time, 8% PFA in H_2_O was directly added to wells 1:1 with wide-orifice tips to minimize cell detachment, and cells were fixed for 25 min at room temperature, after which the sample was treated with the general staining protocol. Extraction times were: 4-5 min (PCNA, MCM2, TIMELESS), 3 min (POLA2, POLD2, POLE2), and 2 min (ND-Cdt1 and MCM2 in MCF-10A SoRa imaging and MCM2 in RPE-1).

#### EdU incorporation and labeling

If measuring 5-ethynyl-2′-deoxyuridine (EdU) incorporation, cells were pulsed with 50 µM EdU (Cayman Chemical Cat# 20518) in growth media for 8 min prior to fixation and pre-extraction, unless otherwise stated. After blocking cells (prior to primary antibodies), cells were washed once with PBS and then a click reaction (Salic and Mitchison, 2008) was performed in 2 mM CuSO_4_, 20 mg/mL sodium ascorbate in TBS (Tris 50 mM, NaCl 150 mM pH 8.3) with 3 µM AFDye 488 picolyl azide (Click Chemistry Tools Cat# 1276) or AFDye 647 picolyl azide (Click Chemistry Tools Cat# 1300) for 30 min, followed by a PBS wash.

#### Antibodies

The following primary antibodies were used for immunofluorescence: rabbit anti-Cdt1 (1:500, Abcam Cat# ab202067, RRID:AB_2651122), rabbit anti-Geminin (1:800, Atlas Antibodies Cat# HPA049977, RRID:AB_2680978), mouse anti-Cyclin A (1:250, Santa Cruz Biotech Cat# sc-271682, RRID:AB_10709300), mouse anti-PCNA (1:200, Santa Cruz Biotech. Cat# sc-56, RRID:AB_628110), rabbit anti-MCM2 (1:800, CST Cat# 3619, RRID:AB_2142137), rabbit anti-p21 (1:2500, CST Cat# 2947, RRID:AB_823586), rabbit anti-HA tag (1:1000, CST Cat# 3724, RRID:AB_1549585), rabbit anti-CDC45 (1:100, CST Cat# 11881, RRID:AB_2715569), rabbit anti-POLA2 (1:100, Atlas Antibodies Cat# HPA037570, RRID:AB_10672280), rabbit anti-POLD2 (1:100, Atlas Antibodies Cat# HPA026745, RRID:AB_1855520), rabbit anti-POLE2 (1:100, Atlas Antibodies Cat# HPA027555, RRID:AB_10610282), rabbit anti-Timeless (1:800, Abcam Cat# ab109512, RRID:AB_10863023), rabbit anti-phospho-Chk1(S317) (1:500, CST Cat# 12302, RRID:AB_2783865), rabbit anti-phospho-Chk2(T68) (1:200, CST Cat# 2661, RRID:AB_331479), rabbit anti-phospho-histone H2A.X(S139) (1:500, CST Cat# 2577, RRID:AB_2118010), rabbit anti-RPA70/RPA1 (1:200, Abcam Cat# ab79398, RRID:AB_1603759). The epitopes for anti-Cdt1 and anti-Geminin antibodies do not detect hCdt1(1-100)^ΔCy^ and hGeminin(1-110) of the CRL4^Cdt2^ and APC/C^Cdh1^ reporters. For secondary antibodies, antibodies targeting the appropriate species and with no spectral overlap were selected from the following and diluted 1:1000: goat anti-rabbit IgG Alexa Fluor 647 (Thermo Cat# A-21245, RRID:AB_2535813), goat anti-rabbit IgG Alexa Fluor 514 (Thermo Cat# A31558, RRID:AB_10375589), goat anti-mouse IgG Alexa Fluor 647 (Thermo Cat# A-21235, RRID:AB_2535804), goat anti-mouse IgG Alexa Fluor 514 (Thermo Cat# A-31555, RRID:AB_2536171)

### Microscopy

#### Time-lapse imaging, RT-QIBC and QIBC

For automated epifluorescence microscopy, cells were imaged using a Ti2-E inverted microscope (Nikon) or ImageXpress Micro XLS microscope (Molecular Devices). For imaging on the Ti2-E, multichannel fluorescent images were taken with triple-band (ECFP/EYFP/mCherry, Chroma: 89006) or quad-band (DAPI/FITC/TRITC/Cy5, Chroma: 89402) Sedat filter sets using an LED light source (Lumencor Spectra X) and Hamamatsu ORCA-Flash4.0 V3 sCMOS camera. A 10x (Nikon CFI Plan Apo Lambda, NA 0.45) or 20x (Nikon CFI Plan Apo Lambda, 0.75 NA) objective was used to acquire images. For imaging on the ImageXpress, images were taken with appropriate single-band filter sets with a white-light source, using a 10x (Nikon CFI Plan Fluor, NA 0.3) or 20x (Nikon CFI Plan Apo Lambda, 0.75 NA) and Andor Zyla 4.2 sCMOS camera. All images were acquired in 16-bit mode without camera-binning, and acquisition settings were chosen to not saturate the signal. Fluorophores and imaging channels were chosen to minimize bleedthrough, and in the case of detectable bleedthrough, it was corrected using bleedthrough coefficients estimated from single fluorophore controls.

For live-cell time-lapse imaging, 96-well plates were imaged within an enclosed 37°C, 5% CO2 environmental chamber in 200 µL of growth media. 4-9 sites were imaged in each well (with the number of wells imaged varying depending on experiment and imaging interval) every 3-12 min. Light exposure to cells was limited by using the minimum exposure necessary to maintain an acceptable signal-to-noise ratio on a per-channel basis, and total light exposure was always limited to below 300 ms per site each timepoint. Images were taken with the 10x objective for all live-cell imaging with the exception of experiments shown in Figure 3A to maximize the number of cells in the field of view. When performing the live-cell imaging for RT-QIBC, cells were immediately taken off the microscope following the final time point and fixed.

For fixed-cell imaging for RT-QIBC and QIBC, tiled images of the majority of each well (16-36 sites per well) were taken using the 20x objective. When reimaging fixed cells (matching back to either live-cell imaging for RT-QIBC or previous rounds of fixed-cell imaging), the plate position (which can shift slightly when replacing the plate on the microscope) was aligned to approximately the same location, and further aligned computationally during image analysis.

#### Spinning-disk confocal microscopy

For live-cell imaging of EYFP-PCNA and Cdt1-mCherry (Figures 1B and S1A), cells were imaged on an automated spinning-disk confocal microscope (Intelligent Imaging Innovations, 3i). This system used a Nikon Ti-E stand, motorized XY stage with piezoelectric Z movement (3i), Andor Zyla 4.2 sCMOS camera, CSU-W1 confocal scanner unit (Yokogawa) and 405/445/488/514/561/640 nm LaserStack (3i), controlled using SlideBook 6 (3i). Cells were imaged in a 37°C environmental chamber (growth media was HEPES buffered), using a 60x/1.35NA oil objective (Nikon) with 2x camera binning. Images at the nucleus midplane were taken every 2-3 min in a 5×5 montage which was stitched together after acquisition. H2B-mTurquoise, EYFP-PCNA and Cdt1-mCherry were imaged using a triple-band 445/515/561 excitation filter set.

For fixed-cell imaging of chromatin-bound ND-Cdt1 localization, pre-extracted MCF-10A cells were imaged on a SoRa spinning-disk confocal microscope (Marianas system, 3i). This system was similar to the previously described 3i microscope, except it used a Zeiss Axio Observer 7 stand, ORCA-Fusion BT sCMOS Camera (Hamamatsu), CSU-W1 SoRa confocal scanner unit (Yokogawa) and 405/445/488/514/561/637 nm LaserStack (3i). Cells were stained using rabbit anti-HA and anti-rabbit Alexa Fluor 488 (for detecting ND-Cdt1) together with mouse anti-PCNA and anti-mouse Alexa Fluor 568. Images were taken using the 488 and 561 channels using a quad-band 405/488/561/640 nm excitation filter set (3i), with a 63x/1.4NA Plan-Apochromat Oil M27 objective (Ziess) and 4x magnification changer and no camera binning. The field of view was manually searched without 4x magnification and low exposure in the 488 channel to identify cells that were positive for ND-Cdt1 expression. Cells were then imaged in both channels at 5 Z-positions around the midplane of the nucleus (0.75 µm spaced, only the midplane is shown). No deconvolution was performed, and controls were tested to ensure there was no spectral bleedthrough or cross-binding of secondary antibodies.

#### Protein nomenclature

For simplicity, we refer to several human proteins by their colloquial names. Namely, we refer to Cdt1 (encoded by *CDT1* gene), Geminin (encoded by *GMNN* gene), Cyclin A (in somatic cells only Cyclin A2, encoded by *CCNA2* gene, is expressed), CRL4^Cdt2^ (Cdt2 is encoded by *DTL* gene), APC/C^Cdh1^ (Cdh1 is encoded by *FZR1* gene), SCF^Skp2^(also known as CRL1^Skp2^, Skp2 is encoded by *SKP2* gene), Cdc6 (encoded by *CDC6* gene), Chk1 (encoded by *CHEK1* gene), Chk2 (encoded by *CHEK2* gene), Wee1 (encoded by *WEE1* gene), p21 (encoded by *CDKN1A* gene), p53 (encoded by *TP53* gene) and Cyclin E (both Cyclin E1 and E2, encoded by *CCNE1* and *CCNE2* genes). Furthermore, we measure several protein complexes through an individual subunit (all of which are constitutive complexes): DNA polymerases epsilon (measured by subunit POLE2), alpha (measured by subunit POLA2) and delta (measured by subunit POLD2), and RPA (measured by subunit RPA1, also known as RPA70).

### Quantification and statistical analysis

#### Image analysis

Automated analysis of time-lapse imaging of cell cycle reporters, quantitative image-based cytometry (QIBC), and Retrospective Time-lapse Synchronized QIBC (RT-QIBC) were performed using a custom MATLAB (R2020a, MathWorks) pipeline based on previous work (Cappell et al., 2016). QIBC here is considered to be the high-throughput single-cell quantification of fixed-cell signals (fluorescent proteins, immunofluorescence, EdU staining, DNA stain), while RT-QIBC involves the assignment of QIBC measurements to an explicit time in the cell cycle based on prior time-lapse imaging of cell cycle reporters (N- and C-CRL4^Cdt2^ reporters, APC/C reporter and EYFP-PCNA). In principle, RT-QIBC can be used to quantify any fixed-cell signal (the techniques used in this study, as well as mRNA or DNA FISH for example) and retrospectively analyze any live-cell reporter or imaging measurements. Image processing pipeline and code used to generate all figures in this study have been deposited on Github and Zenodo (https://github.com/MeyerLab/image-analysis-ratnayeke-2021, DOI: 10.5281/zenodo.5037903), and data can be downloaded at Dryad (https://doi.org/10.5061/dryad.4xgxd2599).

#### Segmentation and signal quantification

Raw images were flat-field corrected (also known as shading corrected) to correct for uneven sample illumination. Since images output by the sCMOS camera are the sum of a camera offset value together with the actual detected signal (which is proportional to the sample illumination), we subtracted off the camera offset value and then divided the image by an empirically determined illumination profile. This profile was calculated either from the background autofluorescence bin areas without cells in live-cell images (aggregated over a large number of sites), or from sample-free wells filled with autofluorescent blocking buffer A for fixed cell imaging. Confocal movies were not flat-field corrected due to a lack of uneven illumination.

For live-cell imaging, nuclei were automatically segmented from H2B-mTurquoise signal using a Laplacian of Gaussian blob detector, which in the case of movies with low contrast, was further refined with active contours. For fixed-cell imaging, nuclei were segmented from the Hoechst stain using a threshold determined from histogram curvature. Detected nuclei larger than the median object size were checked using a curvature-based object splitting algorithm which splits cells along two points of high perimeter-curvature. If there are more than two putative split points, pairs of points are chosen based on pairs with the highest distance along the perimeter between points divided by the Euclidean distance of the points. For multi-round fixed cell imaging, each imaging round was segmented and aligned to each other. Segmentation mask from a single round (typically the first round) was designated the primary mask and used for quantification of all rounds.

To quantify nuclear cell cycle reporters and fixed-cell signals, the background signal was estimated by taking the 25^th^ percentile of pixels outside of a dilated nuclear mask (dilated 7.8 µm for predominantly nuclear signals, 15.6 µm for signals with cytoplasmic component) and subtracted off of images. For chromatin-bound CDC45, POLA2, POLD2, POLE2, and TIMELESS signals, the background was not subtracted during image processing but accounted for later during analysis. The mean and median signal within the nucleus were then calculated, and for signals with a cytoplasmic component, the median signal within a ring outside of the nucleus was calculated (region 0.65 µm to 3.25 µm outside the edge of the nucleus). To quantify puncta area of EYFP-PCNA, a top-hat filter (3 pixel radius for confocal imaging, 2 pixel radius for wide-field) was applied to the image and a series of thresholds of different stringencies were manually chosen and applied to minimize false positives and negatives. The total area of pixels above the thresholds were quantified.

#### Time-lapse tracking

For time-lapse imaging, nuclei were tracked using a nearest-neighbor algorithm between each frame and its previous frame. To increase tracking fidelity, the total nuclear signal (the sum of nuclear intensity) was used as an invariant quantity which does not change significantly over between frames. Using this, putative aberrant merging and splitting of nuclei during segmentation could be detected and corrected. Mitotic events are detected when two daughter nuclei are detected within the vicinity of a previous nucleus and have a total nuclear signal which is approximately equal to the previous nucleus.

To match fixed-cells to live cells tracking, fixed-cell images were computationally aligned to live-cell images using 2D cross-correlation, and cells with their associated measurements were assigned to their nearest live-cell neighbor. When matching the 20x fixed-cell images back to 10x live-cell images, live images were resized using bicubic interpolation (for alignment and tracking purposes only) or fixed images were mean-value binned.

#### Cell cycle annotation of live-cell data

Mitosis was annotated during the process of tracking cells, defined at the separation of the two sets of chromosomes at anaphase. CRL4^Cdt2^ activation (defined as the start of CRL4^Cdt2^ reporter degradation) and Cdt1-mCherry degradation start was annotated by subjecting traces following mitosis or serum-release to a drop detection algorithm. This algorithm detects degradation of the reporter at a given time based on a set number of frames following it (the number of frames after a corresponds to the minimum detectable time since degradation start, typically ∼3 frames). By detecting points using only a set number of frames beyond the degradation point, we avoid biases in the accuracy of detecting cells that just recently degraded compared to cells that degraded much earlier. Points were checked based on the slope and curvature of the trace within the window being low and high enough, respectively, and having a set decrease in the reporter signal (normalized to reporter expression). APC/C^Cdh1^ inactivation (defined as the start of APC/C reporter accumulation) was detected in a complementary way, identifying the first point where the slope increases to a threshold level and the reporter increases from a low level to a threshold-value of persistent increase. All threshold values were empirically determined and validated by eye on at least 200 traces. For both CRL4^Cdt2^ and APC/C reporters, the integrated intensity within the nucleus was quantified for trace analysis.

For identification of the start of S phase from PCNA foci, foci were segmented, and the total area of foci was quantified. The transition from G1 to S phase is characterized by an increase of low foci signal to high foci signal. For RT-QIBC analysis, the same algorithm was used for the APC/C reporter (as the increase in puncta area mirrors the rise in APC/C reporter levels). For confocal imaging, a dual threshold algorithm was used. A high foci signal threshold that robustly identified S phase cells was manually determined (50 pixels), and the first point at which the foci area increased above this threshold and was higher than the previous 4 frames was identified. Since the high foci signal threshold typically identified a point well after S phase entry, the final frame before this high identified point which was below a low threshold (3 pixels) was identified as the true S phase entry point.

#### RT-QIBC

After automated tracking and quantification of live- and fixed-cell imaging, each cell was associated with its corresponding annotated cell cycle reporter traces, as well as multidimensional fixed-cell measurements from QIBC. Based on this, the time elapsed from a point of interest (such as CRL4^Cdt2^ activation or mitosis) was used to arrange fixed-cell measurements based on time. Conversely, live-cell traces can be selected based on QIBC measurements (such as the expression of ND-Cdt1). For analyses with high time resolution (e.g. Figure 1E), time offsets for each imaging site and well were accounted for based on the order of well acquisition.

#### Quantification corrections

For experiments with chromatin-bound proteins measured after pre-extraction, there were rare sections of the samples that were incompletely extracted of soluble proteins. As a proxy for extraction efficiency, in experiments with APC/C or Cyclin E/A-CDK reporters (which are soluble), the residual fluorescent protein signal was imaged in addition to immunofluorescence. Cells that had high fluorescent protein signal for the reporters were considered to be incompletely extracted and removed from the analysis.

For the staining of replication factors in Figures 6 and S6, staining was performed in two rounds. In the first round, chromatin-bound replication proteins (CDC45, TIMELESS, POLE2, POLA2, POLD2, PCNA) were stained using Alexa Fluor 647 secondary antibodies simultaneously with EdU staining with AFDye 488. Fluorophores were then bleached for 1 h, and re-stained for PCNA using an Alexa Fluor 647 secondary antibody to identify S phase cells. However, there was a low-intensity residual signal from the first round of staining, which was corrected for using an empirically determined residual signal scaling factor.

For Figure S3A and S3B, H2B-iRFP670 was expressed in a bicistronic vector together with the APC/C reporter (P2A sequence). As a result, the APC/C reporter signal could be normalized by the H2B-iRFP670 signal to control for differential expression of the construct between cells.

For pre-extraction experiments of replication factors (Figure 6 and S6), outliers resulting from incompletely extracted cells and imaging artifacts were removed by removing the top 1 percentile of data. In Figure S2B, the outer 1 percentile of CRL4^Cdt2^ activation delays from APC/C^Cdh1^ inactivation was removed to account for misidentified cell cycle transitions.

#### Data normalization

For normalized stain quantification, a baseline was calculated from G1 levels of the stain and subtracted off of all values followed by division by the group to be normalized. For EdU quantification shown on a linear scale (for dose-responses and chromatin-bound stain linear fits), the G1 background signal was subtracted off of values for a true zero. For Figure 1F and 1G, measurements were normalized to the median G2 signal of each protein. For Figures 4E and S4E, the EdU signal was zeroed and divided by the Dox(-) EdU signal 1 – 1.2 h after S phase entry to standardize values between replicates.

## Statistical analysis

Details of statistical tests can be found in the figure legends. Generally, comparisons were made with either paired *t*-tests (for tests between multiple independent replicate experiments) or two-sample *t*-tests for within-experiment comparisons of measurements with an α of .05.

For linear fits of chromatin-bound stains (Figure 6), a linear model with a fixed zero-intercept was fit using robust fitting with a bisquare weight function (tuning constant of 2). For fitting dose-response curves, single-cell measurements of EdU and the expression of ND-Cdt1 were fit to a Hill equation of the form

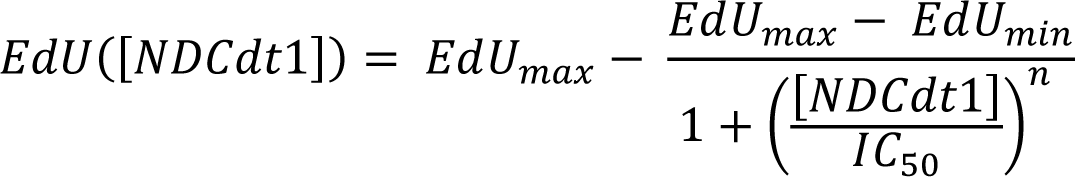

, using nonlinear regression. *EdU_max_* is the EdU incorporation of cells without ND-Cdt1, which was set using cells not expressing ND-Cdt1. *EdU_min_* represents the minimum EdU incorporation, *IC*_50_ is the 50% inhibitory concentration of ND-Cdt1 concentration [*NDCdt*1], and *n* is the Hill coefficient For Figure 3D and 5D, *EdU_min_*, *IC*_50_ and *n* were all fit parameters, while for Figures 3E, 3D, S3D and S3E, *EdU_min_* was set based on high levels of ND-Cdt1 expression. Nonlinear regression was performed using the Levenberg-Marquardt algorithm (nlinfit() in MATLAB). Initialization parameters for *EdU_inhib_* were estimated from the 5^th^ percentile of EdU signal, while *IC*_50_ was initialized based on the median [*NDCdt*1] in the cell population, and *n* was initialized as 1.

For bootstrapped estimators, samples were resampled at least 1000 times and confidence intervals were calculated using the percentile method. For raincloud plots (Allen et al., 2019), which are a combined violin and jitter plot, the data distribution was estimated using a kernel smoothing density estimate. The solid and dashed lines in the violin plot correspond to the median and inter-quartile range (IQR).

To estimate thresholds in an automated manner for determining cells positive and negative for QIBC staining (for example chromatin-bound PCNA positive cells for S phase cells), cells known to be in either G1 or S phase based on live-cell imaging were identified, and then the 99^th^ percentile was chosen as the threshold unless otherwise stated.

### Visualization

For all example cells except for those shown in Figure 7C were extracted from full-sized images through MATLAB scripts and selected based on RT-QIBC or time-lapse analysis. For Figure 7C, cells were selected for imaging based on being in early or late S phase. An example cell was chosen and visualized using ImageJ (v1.53, Fiji distribution) (Schindelin et al., 2012; Schneider et al., 2012) with the QuickFigures plugin (Mazo, 2020).

---

**Figure S1.**
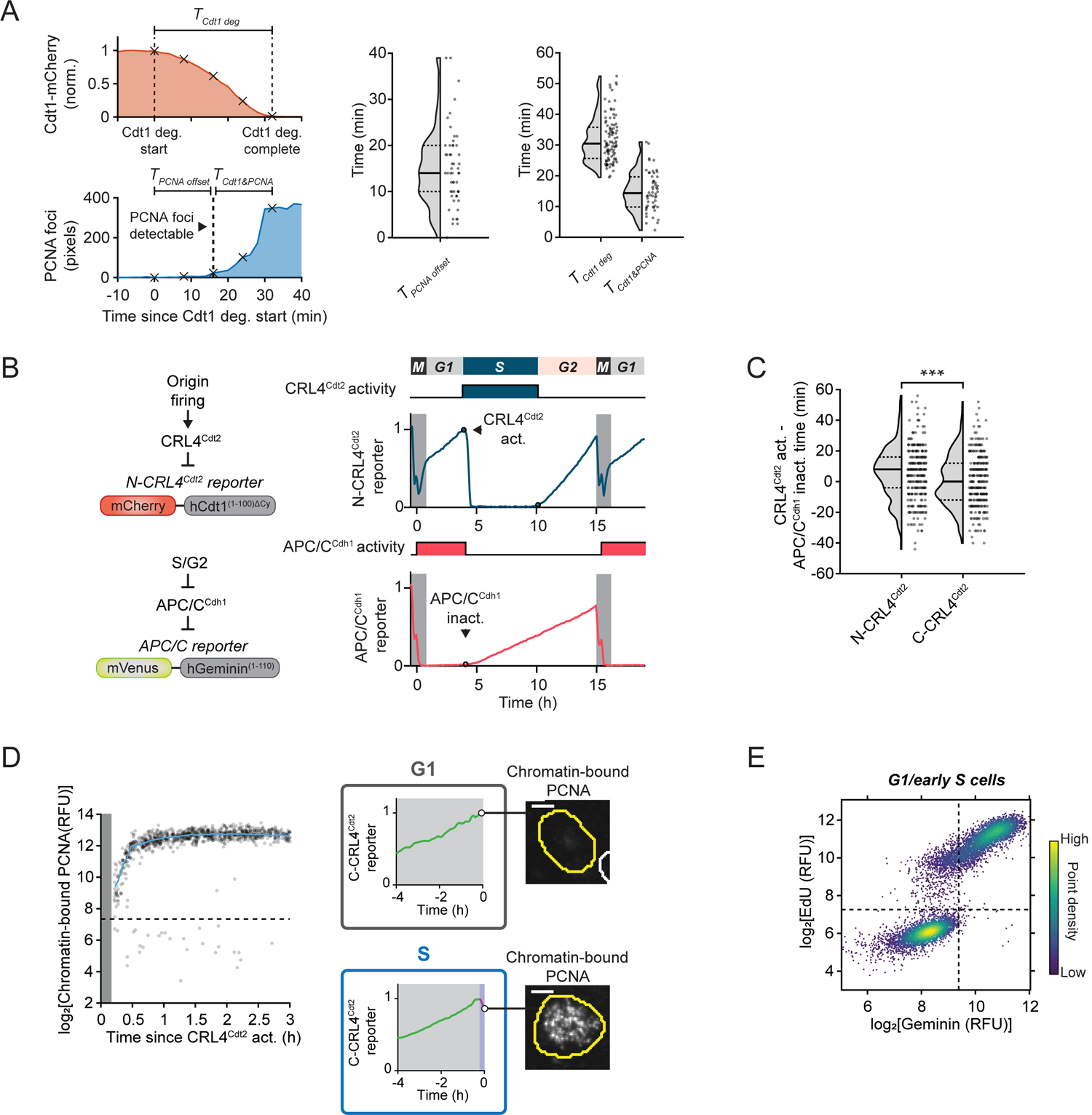
Related to Figure 1 **(A)** MCF-10A cells expressing EYFP-PCNA and doxycycline-inducible Cdt1-mCherry (induced 6 h prior to imaging) were imaged every 2 min using confocal microscopy. Quantification of PCNA foci detection relative to Cdt1 degradation in cells from Figure 1B. Left: Reporters were quantified to determine the time between the start of Cdt1 degradation and first detectable PCNA foci (*T_PCNA offset_*), the total time it takes for Cdt1 to be degraded (*T_Cdt1 deg_*) and the overlap duration when Cdt1 and PCNA foci are simultaneously visible (*T_Cdt1&PCNA_*). Black × corresponds to frames shown in example cell in Figure 1B. Middle: Quantification of the delay in PCNA foci detection from the start of Cdt1 degradation (*T_PCNA offset_*, n = 54 cells). Offset likely represents the amount of time it takes for PCNA foci to grow large enough to be detectable over soluble pool of PCNA. Right: Quantification of *T_Cdt1 deg_* (n = 99 cells) and *T_Cdt1&PCNA_* (n = 54 cells). Cells pooled from 4 independent experiments. Dashed and solid lines in violin plots are IQR and median, respectively. **(B)** Live-cell reporters of CRL4^Cdt2^ and APC/C activity. APC/C reporter is degraded throughout G1 and rises after APC/C^Cdh1^ inactivation. Example traces of N-CRL4^Cdt2^ reporter and APC/C reporter. **(C)** Quantification of CRL4^Cdt2^ activation timing for N-CRL4^Cdt2^ and C-CRL4^Cdt2^ reporters relative to APC/C^Cdh1^ inactivation. Positive values signify CRL4^Cdt2^ activation after APC/C ^Cdh1^ inactivation. n = 300 cells each condition, representative of 2 independent experiments. C-CRL4^Cdt2^ reporter is degraded earlier relative to APC/C ^Cdh1^ inactivation (*** *p*-value = 1.3×10^-4^, two-sample *t*-test, 95% confidence interval 2.6-8.0 min earlier), indicating it is slightly more responsive to initial origin firing. Dashed and solid lines in violin plots are IQR and median, respectively. **(D)** RT-QIBC of chromatin-bound PCNA after degradation of the C-CRL4^Cdt2^ reporter starts. Cells were live-imaged every 3 min, and at the end of imaging, cells were immediately pre-extracted and stained for PCNA. Left: Dashed line is PCNA threshold (95^th^ percentile of G1 cells, 1-2 h after mitosis). Grey bar is time period that is not observed due to the need to have 12 min of reporter degradation to call S phase start. Cells that were identified as having degraded the C-CRL4^Cdt2^ reporter have chromatin-bound PCNA, indicating that origin firing has occurred. The small percentage of cells below the threshold (2.51% of 10,113 cells) had misidentified C-CRL4^Cdt2^ degradation, verified manually. Representative of 2 independent experiments. Right: Example traces and chromatin-bound PCNA stain in G1 (no C-CRL4^Cdt2^ degradation) or just after S phase entry (C-CRL4^Cdt2^ degradation). 5 µm scale bar. **(E)** RT-QIBC in cycling cells of endogenous Geminin and EdU incorporation in cells in G1 to early S (cells selected 3-7 h post mitosis, n = 9,605 cells, representative of 3 independent experiments). Lines demarcate Geminin and EdU positive/negative regions (99^th^ percentile of G1 cells). Note large population of EdU positive, Geminin negative cells, indicating cells which entered S phase with low Geminin.

**Figure S2.**
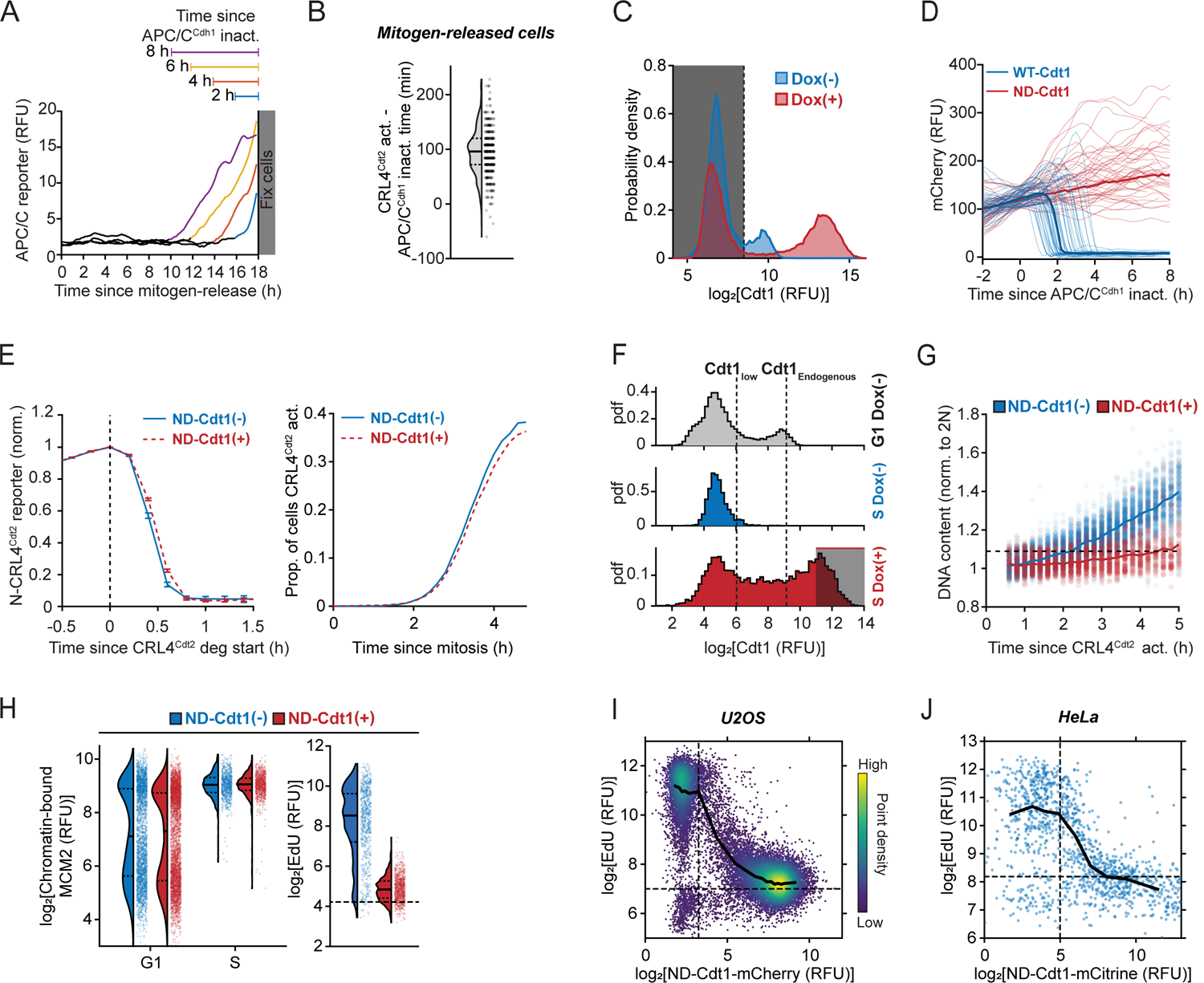
Related to Figure 2 **(A)** Example traces of cells from Figure 2B that inactivated APC/C^Cdh1^ 2, 4, 6 and 8 h prior to fixation. **(B)** Quantification of time between APC/C^Cdh1^ inactivation and CRL4^Cdt2^ activation (N-CRL4^Cdt2^) in mitogen-released cells (compare to cycling cells, Figure S1C). Dashed and solid lines in violin plots are IQR and median, respectively. n = 400 cells, representative of 3 independent experiments. Analyzed from same experiments as Figure 2F. **(C)** Cells from same experiment as Figure 2B. Quantification of Cdt1 immunofluorescence (IF) staining (detecting endogenous Cdt1 as well as Cdt1-mCherry). Mitogen-released cells not expressing Cdt1-mCherry, compared to cells expressing Cdt1-mCherry (doxycycline (Dox)- induced for 24 h). Cells pooled from 5 wells each condition (n ≥ 22,350 cells). **(D)** Quantification of overexpressed Cdt1-mCherry and ND-Cdt1-mCherry as cells enter S phase. Cells induced with Cdt1 constructs with were mitogen-stimulated and live-imaged. n=185 (Cdt1) and 205 (ND-Cdt1) cells. Traces were aligned to APC/C^Cdh1^ inactivation. **(E)** Quantification of N-CRL4^Cdt2^ reporter degradation dynamics with and without ND-Cdt1 induction for 6 h during imaging (in experiment from Figure 2E). Cells were stained for Cdt1 and ND-Cdt1(+) cells were selected in Dox-treated cells. Left: Mean N-CRL4^Cdt2^ reporter intensity following degradation start. Error bars are mean ± 2×SEM (ND-Cdt1(-): n = 547 cells, ND-Cdt1(+): n = 1,644 cells). Right: Proportion of cells entering S phase (N-CRL4^Cdt2^ reporter degraded) over time following mitosis. ND-Cdt1(-): n = 7,637 cells, ND-Cdt1(+): n = 8,880 cells. **(F)** Estimation of ND-Cdt1 expression relative to normal G1 expression levels of Cdt1 measured by IF in mitogen-released cells for Figures 2F, 3D and 3E. Top: Cdt1 in G1 cells without Dox (n = 15,544). Typical endogenous Cdt1 levels at the end of G1 (Cdt1_Endogenous_) was estimated as the median value of cells above the mode of Cdt1 expression in Cdt1 positive cells. Middle: Cdt1 intensity in S phase cells (0.5-1 h after CRL4^Cdt2^ activation) without Dox (n = 250), representing fully degraded Cdt1 levels (Cdt1_low_). Bottom: S phase cells induced with Dox (n = 2,417 cells) in S phase. Shaded bar is gate used for ND-Cdt1(+) cells for Figures 2F. **(G)** Same cells as in Figure 2F examining DNA content measured by Hoechst stain. Values normalized by 2N DNA peak. Line is median value at each time point. ND-Cdt1(-): n = 5,500 cells, ND-Cdt1(+): n = 2,000 cells. **(H)** RT-QIBC of chromatin-bound MCM2 levels (corresponding to licensed origins). ND-Cdt1-mCherry was expressed in mitogen-released cells and the APC/C reporter and ND-Cdt1-mCherry were imaged for 18 h. G1 cells (no APC/C^Cdh1^ inactivation) and S phase cells (1-2 h post APC/C^Cdh1^ inactivation) were identified. Dashed and solid lines in violin plots are IQR and median, respectively. n = 3,418 cells were randomly chosen for each condition shown. Representative of 2 independent experiments. **(I)** QIBC of U2OS cells with ND-Cdt1-mCherry induced by Dox for 6 h. Cells were fixed and Geminin immunofluorescence and EdU incorporation were performed. Early S phase cells were identified as having 2N DNA and positive for Geminin and EdU incorporation, and plotted as a function of ND-Cdt1-mCherry levels. Thresholds were based on G1 EdU signal in cells not induced with Dox. Black line represents median EdU value in bins of ND-Cdt1 levels. n = 19,866 pooled from 8 wells. **(J)** ND-Cdt1-mCitrine was transiently transfected into HeLa cells expressing the APC/C reporter. RT-QIBC was performed after live-imaging both ND-Cdt1-mCitrine and APC/C reporter levels for 15 h. EdU incorporation in early S phase cells (2N DNA, APC/C reporter positive) was measured and plotted according to their ND-Cdt1-mCitrine levels. n = 1,269 cells, pooled from 3 wells.

**Figure S3.**
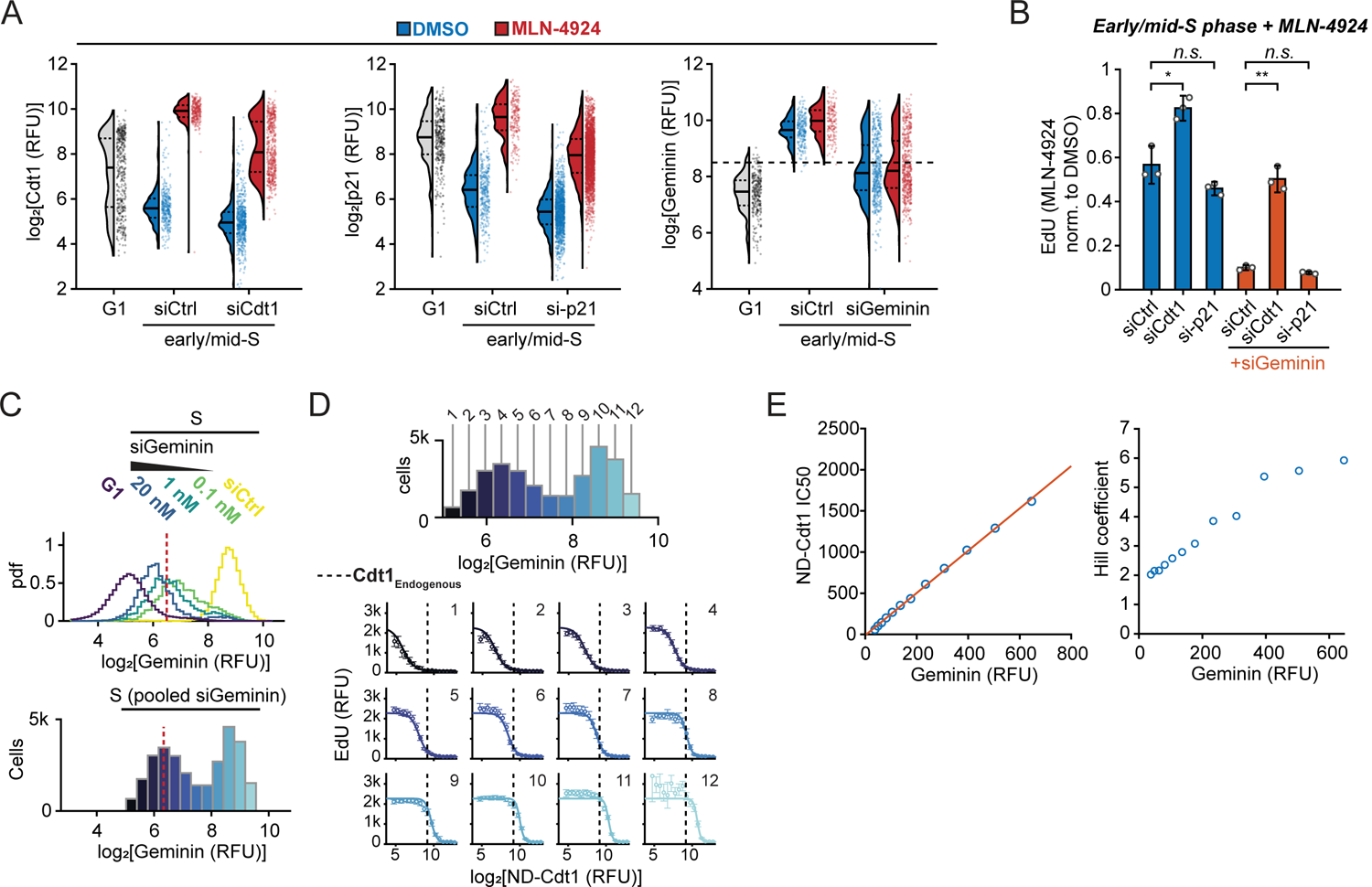
Related to Figure 3 **(A,B)** Cells expressing APC/C reporter and Cyclin E/A-CDK reporter (see Methods and Figure S5A) were serum-starved and treated with siRNA. Cells were then mitogen-released and fixed after 18 h. MLN-4924 was added 4 h prior to fixation. QIBC was performed to quantify the levels of the fluorescent reporters, immunofluorescence and EdU. Gating for early/mid-S phase cells is based on APC/C reporter intensity in 2N-3N cells with high Cyclin E/A-CDK activity ( ≥ 0.8). Data pooled from 3 wells. **(A)** Validation of changes in protein levels with MLN-4924 and siRNA knockdowns. G1 cells were chosen based on being negative for APC/C reporter and EdU incorporation, and intermediate Cyclin E/A-CDK activity (0.5 - 0.8). Dashed and solid lines in violin plots are IQR and median, respectively. n ≥ 174 cells for all conditions. G1 cells were treated with control siRNA to identify normal G1 levels. For right panel, dashed line is threshold below which cells with siGeminin were considered fully knocked down in Figure S3B. **(B)** Impact of siRNA knockdown on EdU incorporation in the presence of MLN-4924 in early/mid-S phase. Points are median values of cells in different wells and bars are mean ± 2×SEM of the medians. n ≥ 57 cells per well. For siGeminin conditions, cells with low Geminin were selected. Two-sample *t*-test: siCtrl – siCdt1(* *p*-value = 3.7×10^-2^), siCtrl – si-p21 (n.s. *p*-value = 8.6 ×10^-2^), siCtrl/siGeminin – siCdt1/siGeminin (** *p*-value = 4.5×10^-3^), siCtrl/siGeminin – si-p21/ siGeminin (n.s. *p*-value = 8.0×10^-2^). **(C)** Top: A range of Geminin levels in cells 2-3 h after S phase entry (N-CRL4^Cdt2^ reporter) were produced by titrating siGeminin (n ≥ 4,572 cells for all conditions) for experiments in Figures 3E and 3F. Dashed line is threshold for low Geminin levels, determined from G1 cell Geminin levels. Representative of 3 independent experiments. Bottom: Pooled S phase cells from all siGeminin conditions to generate a range of Geminin expression. **(D)** Top: Cells from Figure S3C for all siRNA conditions were pooled together and separated into 12 bins for analysis of the impact of Geminin on EdU incorporation. (n = 29,350 total cells). Individual dose-response fits from Figure 3E, same experiment as in Figure 2F. Dashed line represents endogenous Cdt1 levels, determined from Figure S2F. Points and error bars are mean ± 2×SEM for bins of ND-Cdt1 expression for given Geminin level (bins ≥ 6 cells, median bin count 127). **(E)** ND-Cdt1 IC_50_(left) and Hill coefficient (right) for fit dose-response as a function of Geminin expression levels. Left: Line is linear regression fit (R^2^ = .999).

**Figure S4.**
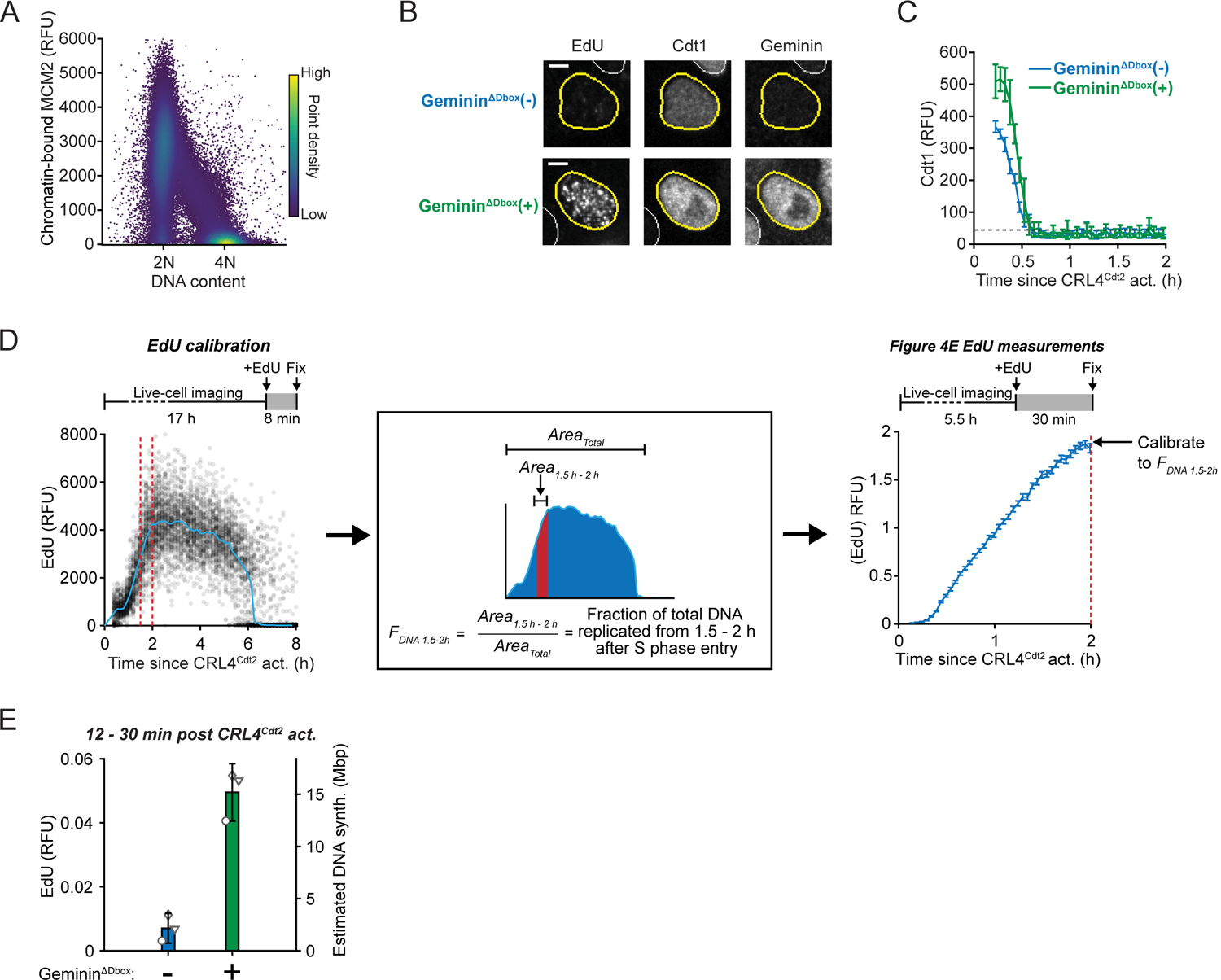
Related to Figure 4 **(A)** QIBC of chromatin-bound MCM2 (n = 96,297 cells) in cycling cells as a function of DNA content. 2N DNA content was estimated from G1 DNA intensity. RT-QIBC analysis from Figure 4D was performed simultaneously with this experiment. **(B)** Same example cells as shown in Figure 4E, co-stained for EdU incorporation, Cdt1 and Geminin (detecting both endogenous Geminin and Geminin^ΔDbox^). Scale bar = 5 µm. Note, Cdt1 levels are likely increased in response to Geminin^ΔDbox^ due to co-stabilization. However, Cdt1 is inactivated by Geminin, and Cdt1 is still degraded over the same 30 min period (Figure S4C). **(C)** RT-QIBC of endogenous Cdt1 following C-CRL4^Cdt2^ activation in cells with Geminin^ΔDbox^ overexpressed. Same cells as in Figure 4E and S4B. Cells that had doxycycline (Dox) added ≤1 h prior to mitosis were analyzed. Geminin^ΔDbox^ does not impact the time it takes to degrade Cdt1. Error bars and line are mean ± 2×SEM in bins of cells (Geminin^ΔDbox^(-): n = 3,436, Geminin^ΔDbox^(+): 2,302 cells total, n ≥ 32 cells per bin). Representative for 3 independent experiments. **(D,E)** Calibration of EdU incorporation to absolute DNA synthesis from Figure 4E using RT-QIBC (C-CRL4^Cdt2^ reporter). In general, the relationship between EdU incorporation intensity and absolute DNA synthesis (in base pairs) can be inferred by integrating EdU incorporation measurements made throughout S phase, which estimates the signal which would be observed if EdU was incorporated in all of S phase. From this analysis, the fraction of total DNA synthesis during a given period (in this experiment, a period 1.5-2 h after S phase entry was chosen as a calibration point. Denoted as F_DNA 1.5-2h_) can be estimated by taking the ratio of the area from 1.5-2 h after S phase entry (Area_1.5-2h_) to the total area (Area_Total_). In the experiment from Figure 4E, a 30 min EdU pulse was used, and thus the EdU intensity in cells 2 h after S phase entry in cells without Dox added would be equivalent to F_DNA 1.5-2h_. **(D)** Left: F_DNA 1.5-2h_ was estimated in cells by RT-QIBC of an 8 min EdU pulse at the end of imaging, with 8 min time interval for live-cell imaging. The median EdU incorporation for each timepoint was calculated and used to determine area under the curve (n = 13,626 cells). Middle: Calculation of area under curve. Left: RT-QIBC measurements of EdU incorporation from a 30 min EdU pulse at the end of imaging. Line and error bars are mean ± 2×SEM in cells within bins. Data pooled from 3 independent experiments (n = 33,208 cells). **(E)** The EdU signal in each condition from Figure 4E was calibrated based on Figure S4D to find the equivalent fraction of total DNA synthesis. Multiplying this by 6×10^9^ base pairs (approximate human diploid DNA) gives the equivalent amount of DNA synthesis in base pairs. For each of 3 independent experiments, the median of cells were taken (n ≥ 31 cells per replicate per condition, Geminin^ΔDbox^(-) 120 cells total, Geminin^ΔDbox^(+) 213 cells total). Error bars are mean ± 2×SEM.

**Figure S5.**
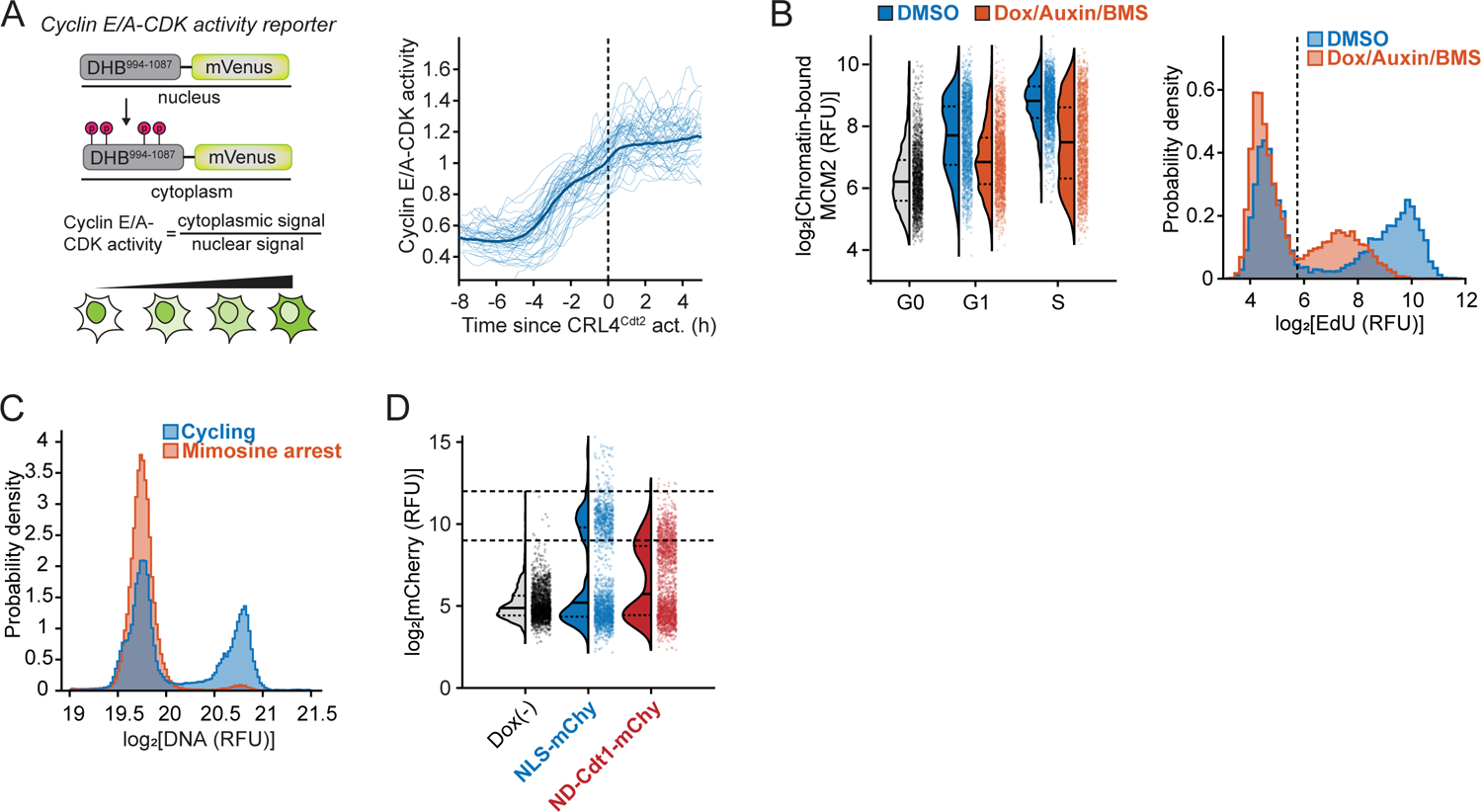
Related to Figure 5 **(A)** Cyclin E/A-CDK activity reporter is initially nuclear localized in G0 and early G1 and gradually translocates to the cytoplasm in response to Cyclin E/A-CDK activity. 50 sample traces in cells released from serum starvation aligned to N-CRL4^Cdt2^ reporter degradation. Thick line is median trace of 1,269 cells. **(B)** RPE-1 *p53^-/-^ CDC6^d/d^* cells were serum-starved in G0. Cells were then mitogen-released in the presence of doxycycline (Dox), Auxin and BMS-650032 (BMS) to degrade Cdc6 as cells re-enter the cell cycle and inhibit origin licensing, or with vehicle (DMSO) to permit origin licensing. Left: Cells were fixed 12 h after serum release and QIBC was performed on chromatin-bound MCM2. G0 cells are unreleased cells, and G1 and early S phase cells were chosen on the basis of EdU incorporation. n = 2,000 cells for all conditions. Pooled from 2 wells in each condition. Right: Cells were fixed 15 h after serum release and QIBC was performed on EdU incorporation. 2N DNA (G1/early S phase cells) were plotted, and dashed line is threshold for EdU incorporation, calculated from G0 cells. n = 12,380 cells (DMSO), 13,543 cells (Dox/Auxin/BMS). Pooled from 2 wells. **(C)** QIBC of DNA content in mimosine arrested cells using protocol in Figure 5F, compared to cycling cells. N = 52,433 cells (mimosine arrested) and 61,472 cells (cycling). Representative of 2 independent experiments. **(D)** QIBC of mCherry fluorescence in cells induced in Figure 5F. n = 2,000 cells per condition. Cells with mCherry fluorescence within lines were chosen for analysis in Figure 5G.

**Figure S6.**
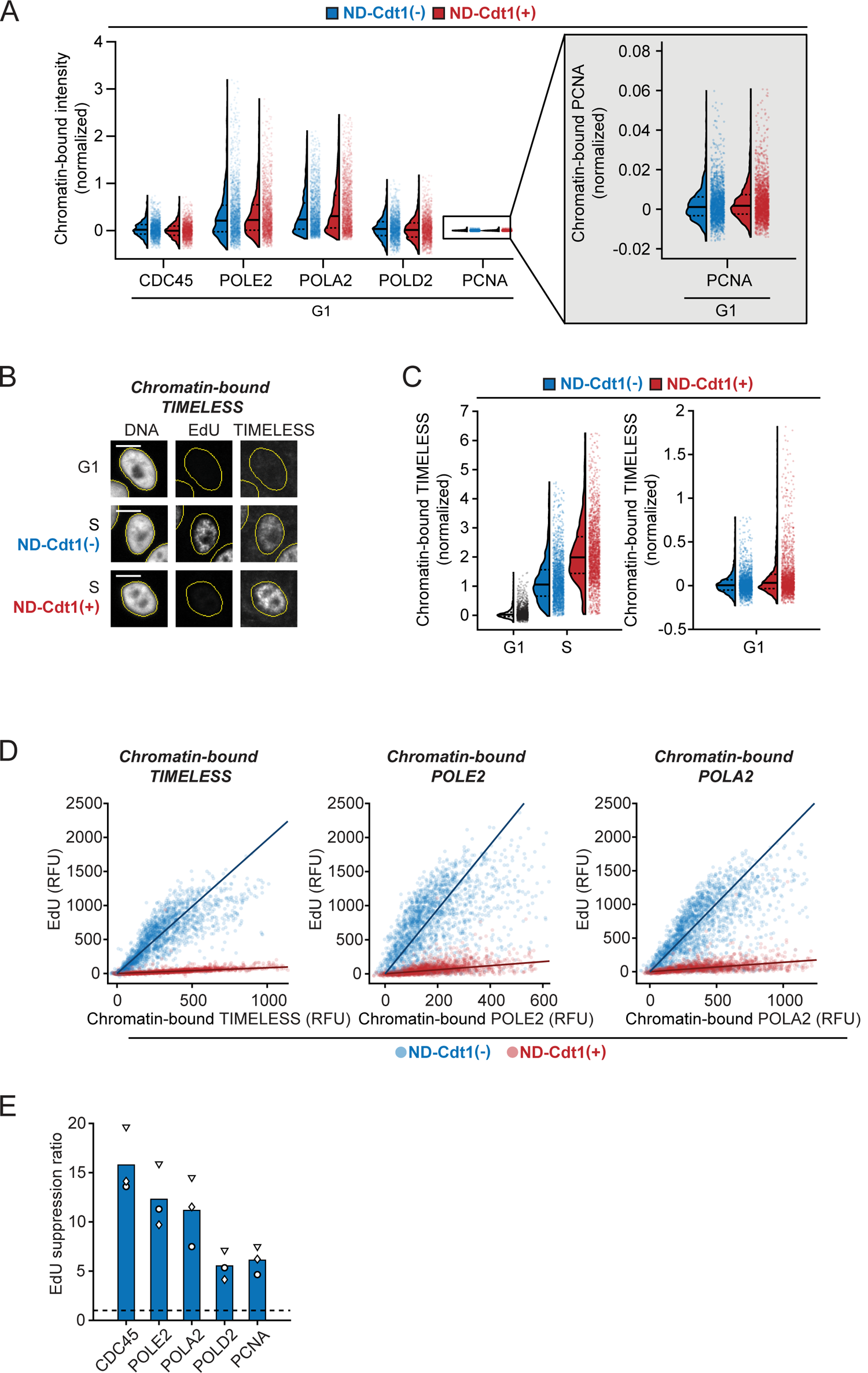
Related to Figure 6 **(A-E)** RT-QIBC of chromatin-bound replisome components in cells expressing ND-Cdt1-mCherry. Mitogen-released cells with the APC/C reporter were induced with doxycycline and released for 18 h. G1 cells had active APC/C^Cdh1^ without chromatin-bound PCNA. Cells which expressed ND-Cdt1-mCherry during live imaging were selected for ND-Cdt1(+). Representative of 3 independent experiments. **(A)** Comparison of chromatin-bound replisome components in G1 with ND-Cdt1, in same experiment as Figure 6B. G1 mode intensities from ND-Cdt1(-) were subtracted off signals and values were normalized to the ND-Cdt1(-) S phase condition from Figure 6B. Dashed and solid lines in violin plots are IQR and median, respectively. N = 2,000 cells per condition. Representative of 3 individual experiments. **(B)** Representative cells of chromatin-bound TIMELESS. Scale bar = 10 µm. **(C)** Comparison of chromatin-bound TIMELESS in S phase (left) and G1 phase (right), analyzed in same way as Figure 6B and Figure S6A. n = 2,000 cells, pooled from 3 wells for ND-Cdt1(-), or 7 wells for ND-Cdt1(+). **(D)** Analysis of EdU incorporation as a function of chromatin-bound protein levels for TIMELESS, POLE2, POLA2 in S phase. G1 mode intensities were subtracted off EdU and chromatin-bound intensity. Line is fit line of linear regression (n = 2000 cells). Representative of 3 independent experiments. Other stains in Figure 6C. TIMELESS staining was done in separate experiment as other stains from this figure and Figure 6C. Cells were pooled from 3 wells for ND-Cdt1(-), or 7 wells for ND-Cdt1(+) from a single experiment. **(E)** Summary of slopes from fit lines from Figure 6C and S6D. EdU suppression ratio is defined as the ratio of the fit line in the control condition to the ND-Cdt1 condition (>1 indicates there is lower EdU incorporation for a given amount of chromatin-bound protein). Bar is mean of 3 independent experiments.

**Figure S7.**
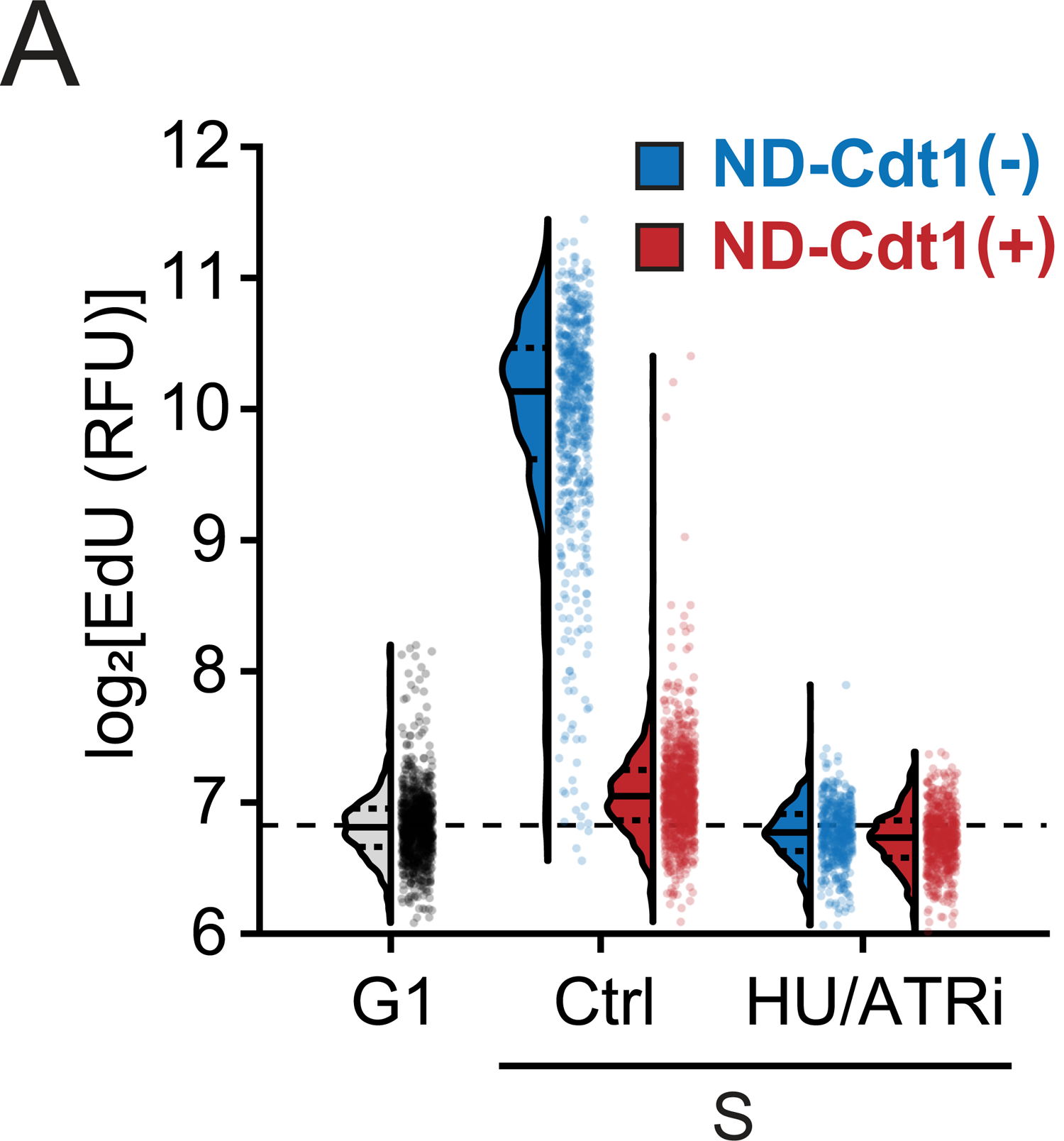
Related to Figure 7 **(A)** RT-QIBC of EdU incorporation in cells treated and analyzed as in Figure 7B (co-stained together with CDC45). n ≥ 503 cells for all conditions. Representative of 3 independent experiments.

**Table S1. Related to STAR Methods** Supplemental details on recombinant DNA used to generate cell lines in this study (and which Figures each cell line was used in), as well as siRNA oligonucleotides used in this study.

